# The effect of a temperature-sensitive prophage on the evolution of virulence in an opportunistic bacterial pathogen

**DOI:** 10.1101/850248

**Authors:** Matthieu Bruneaux, Roghaieh Ashrafi, Ilkka Kronholm, Elina Laanto, Anni-Maria Örmälä-Odegrip, Juan A. Galarza, Chen Zihan, Mruthyunjay Kubendran Sumathi, Tarmo Ketola

## Abstract

Viruses are key actors of ecosystems and have major impacts on global biogeochemical cycles. Prophages deserve particular attention as they are ubiquitous in bacterial genomes and can enter a lytic cycle when triggered by environmental conditions. We explored how temperature affects the interactions between prophages and other biological levels by using an opportunistic pathogen, the bacterium *Serratia marcescens*, that harbours several prophages and that had undergone an evolution experiment under several temperature regimes. We found that the release of one of the prophages was temperature-sensitive and malleable to evolutionary changes. We further discovered that the virulence of the bacterium in an insect model also evolved and was positively correlated with phage release rates. We determined through analysis of genetic and epigenetic data that changes in the outer cell wall structure possibly explain this phenomenon. We hypothezise that the temperature-dependent phage release rate acted as a selection pressure on *S. marcescens* and that it resulted in modified bacterial virulence in the insect host. Our study system illustrates how viruses can mediate the influence of abiotic environmental changes to other biological levels and thus be involved in ecosystem feedback loops.

## Introduction

Viruses in general, and bacteriophages in particular, are increasingly being recognized as key actors of global biogeochemical cycles (Suttle, 2007; Weitz et al., 2015; Shelford and Suttle, 2018). They interact actively with the global climate change via multiple feedback loops (Danovaro et al., 2011). Among bacteriophages, many temperate phages can integrate into the genome of their host as prophages instead of pursuing a lytic life cycle. Such prophages are ubiquitous in bacteria (Touchon et al., 2016; Tuttle and Buchan, 2020) and they constitute a tight evolutionary unit with their host, ranging on a spectrum from mutualistic parasites to domesticated phages, that affects the evolution of bacterial genomes. Understanding the relations between prophages and their microbial hosts and how shifts in their interactions are transmitted and amplified in downstream processes is crucial (Zarnetske et al., 2012; Buck and Ripple, 2017). It is well demonstrated that prophages can impact their hosts and higher biological levels directly (Argov et al., 2017), for example by altering bacterial gene regulation and physiology (Laumay et al., 2019; Schroven et al., 2021), providing virulence factors to their host (Fortier and Sekulovic, 2013), or modifying bacterial abundance in ecosystems when switching to a lytic cycle (Williamson et al., 2002; Brum et al., 2016; Howard-Varona et al., 2017).

Here, we report evidence suggesting that environmentally-inducible prophages can act not only directly but also indirectly as selection pressure “transmission belts” between environmental changes and the evolution of virulence of their bacterial hosts. We used clones from an evolution experiment where the bacterium *Serratia marcescens* had been exposed to three different temperature regimes (Ketola et al. 2013; Figure 1). *Serratia marcescens* is an opportunistic pathogen found in water and soil that can be virulent in plants, insects, and vertebrates, and is responsible for nosocomial infections in humans (Flyg et al., 1980; Grimont and Grimont, 2006). Based on single-molecule real-time (SMRT) sequencing data of experimentally evolved clones (Bruneaux et al. 2021; Figure 1), we showed that those strains contained several prophage candidates and we designed a qPCR-based method to quantify the abundance of extracellular phage particles in liquid cultures. We showed that one of those prophages was released in a temperature-sensitive manner and that phage release rates diverged during the evolution experiment, with strains evolving in prophage-inducing temperature evolving decreased release rates while strains evolving in non prophage-inducing temperature evolving increased release rates. We also found that evolutionary changes in phage release rates were correlated with changes in bacterial virulence in an insect model. While we showed in a previous study that temperature, the environmental selective pressure imposed on bacterial populations during the evolution experiment, was not strongly associated with genetic or epigenetic changes (Bruneaux et al., 2021), we found here that many genetic and epigenetic differences observed between bacteria genomes can be interpreted as prophage-related, and in particular linked to the outer structure of the bacterial cell wall which is itself known to play a role in bacterial virulence. By shedding light on another way that bacteriophages can affect the evolution of their hosts and indirectly impact other biological levels, our results strengthen the understanding of the feedback loops between viruses, ecosystems, and environmental changes.

**Figure 1:**
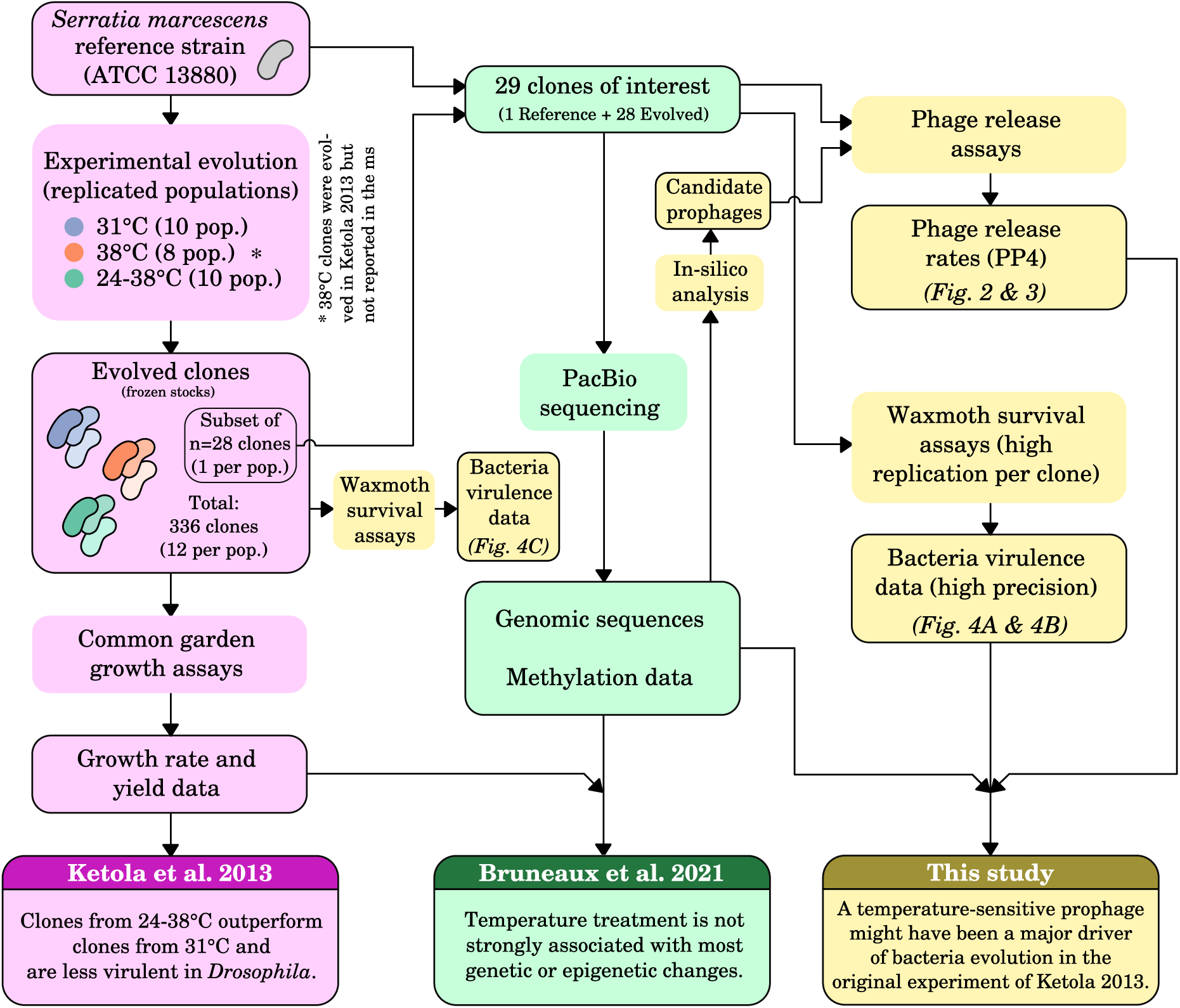
Overview of the relationships between our study and previous studies based on the same evolution experiment (Ketola et al., 2013; Bruneaux et al., 2021). Boxes are color-coded based on which study each item originates from.

## Materials and Methods

### Origin of the strains used in this study

Strains originated from an earlier evolution experiment (Figure 1; Ketola et al. 2013), of which a brief overview is given here. To initiate the experiment, a freshly isolated *Serratia marcescens* ancestor derived from the ATCC 13880 reference strain was grown overnight at 31 °C in low-nutrient medium (SPL 1%, hay extract) and spread to 30 replicate populations (10 populations per treatment). The 400 μL populations were placed under constant 31 °C, constant 38 °C, or daily fluctuating (24-38 °C) thermal treatments, with daily renewal of low-nutrient medium. After evolving for 30 days, 12 clones were isolated from each population by dilution plating on LB agar plates. Clones were grown overnight in low-nutrient medium and frozen to 100-well Bioscreen plates (1:1 with 80 % glycerol) in randomized order.

### DNA extraction, sequencing, and genome annotation

Detailed methods for DNA extraction, PacBio SMRT sequencing, and genome annotation are reported in a previous study (Figure 1; Bruneaux et al. 2021) but a brief overview is given here. One frozen clone per replicate population was randomly chosen for sequencing, as well as the original reference strain. Frozen stocks were grown in overnight precultures followed by 24 h growth in 150 ml of SPL 1%. Since two populations from the constant 38 °C treatment were lost during the experimental evolution, we sequenced 10 clones from constant 31 °C, 10 clones from fluctuating 24-38 °C, and 8 clones from constant 38 °C (Supplementary Figure S1). Bacterial DNA was extracted from pelleted cells and sequenced on a PacBio RS II sequencing platform using P6-C4 chemistry, with two single-molecule real-time sequencing (SMRT) cells run per DNA sample. Standard PacBio analysis protocols were used to assemble genomes and to estimate methylation fractions for adenine bases. Genome annotation was performed using the NCBI Prokaryotic Genome Annotation Pipeline (PGAP; Tatusova et al. 2016).

### Identification of prophages and quantification of release rates

#### Detection of prophage candidates

We used the PHASTER prediction tool (Arndt et al., 2016) (https://phaster.ca/) to detect the presence of prophage candidates in the genome of the *Serratia marescens* reference strain we sequenced. The submission to the PHASTER server was done on 2019-04-21 and five putative prophages were detected (Supplementary Table S1).

#### Phage particle count using qPCR

We developed a method based on quantitative polymerase chain reaction (qPCR) to quantify the amount of extracellular phage particles in liquid cultures of *S. marcescens*. The detailed method and the associated statistical analysis are fully described in the Supplementary Methods, but an overview is given here. We used five specific primer pairs targetting each of the candidate prophage regions and one additional primer pair targetting a chromosomal, non-prophage-related bacterial gene (Supplementary Table S2) to quantify the amount of prophage DNA copies relative to the amount of bacterial genome copies present in a sample using qPCR. In essence, an excess of prophage DNA copies would indicate that phage genomes were replicated outside the bacterial chromosome. To quantify released phage particles in particular, and given low expected induction rates and the logarithmic scale of the uncertainty of qPCR estimates, our approach to reliably quantify the minute excess of extracellular prophage DNA was the following (Supplementary Figure S2): (i) split a bacterial culture sample to be analysed into one raw sample and one supernatant sample obtained after gentle centrifugation to pellet bacteria cells, (ii) process both samples by DNase to digest DNA fragments which were not protected inside a bacterial cell nor inside a phage particle, (iii) inactivate DNase and release DNA from cells and phage particles by heating the samples at 95 °C and (iv) quantify the amount of bacterial genome copies and of prophage DNA copies in both samples with qPCR. The supernatant sample is expected to be impoverished in bacterial cells, while phage particles can remain in suspension, and thus the proportion of prophage DNA copies which were not contained in bacteria cells in the culture (i.e. which were presumably in phage particles) can be estimated from the differential decrease in qPCR estimates for prophage DNA copies and bacterial genome copies between the raw and supernatant samples. A Bayesian model was implemented to estimate phage-to-bacteria ratios based on qPCR results.

#### Quantification of phage release rates in evolved strains

Using the approach outlined above, release rates of the five candidate prophages (i.e. all prophage candidates identified by PHASTER, irrespective of the prediction for prophage completeness) were tested under five temperature assay conditions in the 28 evolved clones selected for sequencing and in the original reference strain. The assays lasted two days, and were made in SPL 1 % under one of the following treatments: 31-31 °C, 24-24 °C, 38-38 °C, 24-38 °C and 38-24 °C, where the temperatures are the temperatures on the first and second day, respectively, with a transfer to fresh medium between them (Supplementary Figure S2). The details of the Bayesian model used to estimate the effect of evolutionary treatment and of assay temperature are given in Supplementary Methods.

### Comparative genomics of prophage PP4

#### Annotation of PP4

Since the candidate prophage PP4 was the only one for which induction was detected in our assays, we focused our annotation and comparative analysis efforts on this prophage. The PHASTER output for PP4 contained a set of contiguous CDS in the *S. marcescens* genome which were related to phage functions. It also provided putative locations of attL/attR attachment sequences but those were located between prophage CDS instead of being at their periphery. We searched for better attL/attR candidates by looking for the longest repeated motif located 3000 bp upstream and downstream of the proposed prophage CDS. We found better candidates for attL/attR which encompassed all the PP4 CDS: the corresponding motif was 20-bp long, compared to the 12-bp long motif found by PHASTER. We defined PP4 as the sequence encompassed by those attL/attR sequences.

Annotation of PP4 CDS was manually curated by merging the annotation obtained for the *S. marcescens* genome (as described above and in Bruneaux et al. 2021) and the annotation provided by PHASTER. A last attempt to identify the CDS for which the “hypothetical protein” status remained at this stage was performed by blasting their predicted protein sequences against the nr database using NCBI blastx server and its default settings, but excluding the *Serratia* taxid (blast run on 2020-04-10).

#### Identification of phages and prophages related to PP4

To identify phages related to PP4, we retrieved all the phage genomes available from RefSeq Nucleotides using the query `Viruses[Organism] AND srcdb refseq[PROP] NOT wgs[PROP] NOT cellular organisms[ORGN] NOT AC 000001:AC 999999[PACC] AND (“vhost bacteria”[Filter])`. The database was searched on 2020-04-01 and returned 2522 phage genomes. The nucleotide sequence of PP4 was compared to those phage genomes using a local blastn search (task blastn). The three phage genomes giving the total best scores were temperate phages: P88 which infects *Escherichia coli* (KP063541) (Chen et al., 2017) and SEN4 and SEN5 which infect *Salmonella enterica* (KT630645 and KT630646) (Mikalová et al., 2017).

To identify bacterial genomes containing prophages related to PP4, we ran a blastn search of the nucleotide sequence of PP4 against the NCBI nt database (excluding *Serratia* taxid) on 2020-04-02. The bacteria genomes giving the best total scores were *Klebsiella oxytoca* KONIH1 (CP008788.1) and *K. oxytoca* KONIH4 (CP026269.1), which gave very similar scores. We selected KONIH1 for further analysis. To determine the precise locations of PP4-related prophages in the chromosome of KONIH1, we ran a local tblastx search of its sequence against PP4 and produced a dot-plot using the tblastx matches. We visually identified three segments showing a consistent matching structure with PP4. For each segment, we determined the best attL/attR candidates using the same approach used when refining the attL/attR sites for PP4 as described above. This enabled us to extract three prophages (KONIH1/1-3). For each of these prophages, we extracted the CDS contained between their attL/attR based on their GenBank record. Finally, we also searched the genome of the *Serratia* strain we used in our study for other prophages similar to PP4, using the same local tblastx approach as described for KONIH1. The dot plot showed that PP7 was related to PP4, and we included PP7 in our final comparisons.

Once we extracted the nucleotide sequences for phages P88, SEN4, KSP20 (two fragments available: AB452988.1 and AB452989.1) and for prophages KONIH1/1-3 and PP7, we identified the matches between their predicted proteins and those from PP4 by running a final local tblastx search of each of those sequences against the PP4 sequence.

### Detection of phage particles by other methods

#### Transmission electron microscopy

Two precultures of the *S. marcescens* reference strain were grown overnight at 31 °C in 10 ml of SPL 1 % and used to inoculate two volumes of 100 ml of SPL 1 %. After 10 hours, the 100 ml cultures were each added to 300 ml of fresh medium and culture continued for 17 hours at 31 °C. Cells from the two 400 ml batches were pelleted separately (20 min at 8000 rpm with Sorvall RC-6+ centrifuge and F12-6x500 rotor) and their supernatants were filtered with 0.45 μm filters. The filtered supernatant from one 400 ml batch was then pelleted (1.5 hour at 24 000 rpm at 4 °C with Beckman Coulter L-90K centrifuge and 45 Ti rotor) and supernatant was discarded. The filtered supernatant from the second 400 ml batch was added, and a second centrifugation (1.5 hour at 24 000 rpm) was applied. Pellets were resuspended in 1 ml of 0.1 m ammonium acetate and combined. After a last round of centrifugation with the same settings, the final pellet was resuspended in a total of 100 μl of 0.2 m Tris-HCl and stored at 4 °C until staining for TEM. For staining, 1:10 dilutions of pellet (10 μl) were placed for 1 minute on glow discharge-treated copper grids. Staining was done with 1 % phosphotungstic acid (PTA) for 2 minutes. Imaging was performed with JEOL JEM-1400HC at 80 kV. The resuspended pellet was also used in qPCR reactions similar to the ones used to quantify prophage induction, using all the candidate prophage-specific primers, to confirm the identity of the phage particles observed in TEM.

#### Plaque assays

The *S. marcescens* reference strain was grown in liquid medium at room temperature and at 37 °C. Supernatants and cells were prepared from both growth conditions. Bacteria (100 μl) were spread on top of 1 % agar plates and the double layer agar method was also employed by mixing 300 μl with 3 ml of 0.7 % soft agar and poured on top of the solid agar. Supernatants (10 μl drops) were applied on plate cultures of the *S. marcescens* reference strain itself, of two freshwater *Serratia* strains, and of Db10 and Db11 *Serratia* strains grown at room temperature and at 37 °C. No plaque was observed for any combination of growth condition of the reference strain with any receiving plate. Additionally, a separate experiment where mitomycin C was added (1 μg ml*^−^*^1^) to liquid cultures of the *S. marcescens* reference strain either at room temperature or at 37 °C did not result in any visible clearing.

### Bacterial virulence assays in an insect model

We estimated the virulence of our *S. marcescens* strains by measuring the longevity of waxmoth larvae (*Galleria mellonella*) injected with 5 μl of bacterial culture. *G. mellonella* is an interesting model for bacterial virulence as it had been shown that virulence in this host is also a good proxy for virulence in vertebrate hosts such as mouse or fish (Jander et al., 2000; Djainal et al., 2020). We grew bacterial cultures of evolved strains overnight at 31 °C in Bioscreen wells in 400 μl of SPL 1 % inoculated with the strains frozen stocks using a cryoreplicator. The reference strain was similarly grown overnight at 31 °C in 8 ml of SPL 1 % in a loose-capped 15 ml tube inoculated from a frozen sample. On injection day, culture optical densities were measured and each larva was injected with 5 μl of a single culture in the hemocoel with a Hamilton syringe. For each strain, 20 larvae were injected simultaneously; ten of those were then incubated at 24 °C while the other ten were incubated at 31 °C. Larval survival was monitored at 1-3 hour intervals by checking for larva movements, and time of death was recorded as the inspection time when a larva was found unresponsive. Additionally, for each incubation temperature, ten larvae were injected with sterile medium and ten with sterile water as controls. This setup was replicated four times, resulting in a total of 80 infected larvae per strain (40 to incubation at 24 °C and 40 to incubation at 31 °C). Some larvae from the first replication block were discarded due to a technical problem, leaving three replication blocks instead of four for some strains.

We analysed the larval survival data using a Cox proportional hazards model, where replication block, larval body mass, culture optical density, strain identity, incubation temperature and the interaction between strain identity and incubation temperature were included as fixed effects. In this type of model, the hazard function describes the instantaneous rate of death at a given time *t* for an individual still alive at *t*. The model included the effect of strain evolutionary treatment on their virulence, using a hierarchical Bayesian approach in JAGS 4.1.0 (Plummer et al., 2003; Su and Yajima, 2015) with the R2jags package. The proportional hazards were implemented as described by Clayton 1991 (Clayton, 1991) based on code from the OpenBUGS Examples (The OpenBUGS Project). The details of the model are presented in the Supplementary Methods.

### Analysis of genetic and epigenetic variation

We used the genomic and methylation data for the 28 evolved clones of interest and the reference strain available from Bruneaux et al. 2021 (Figure 1). As reported in this previous study, no evidence of genetic polymorphism was found within each sequenced sample (i.e. each sequenced clone was indeed genetically clonal) and loci variable across clones were identified from an alignment of the 29 sequenced genomes produced by Mugsy (Angiuoli and Salzberg 2011; Supplementary Table S3). To investigate the association between genetic variation and phenotypic traits (phage release and virulence in waxmoth larvae), we ran Wilcoxon rank sum tests for all combinations of genetic variants and traits, using only genetic variants present in at least two strains. P-values were corrected for multiple testing using the false-discovery rate method (Benjamini and Hochberg, 1995).

Epigenetic data consisted of the methylation fraction for adenosine bases in all GATC motifs present in the reference strain genome (38 150 GATC palindromes were present in the reference strain genome, corresponding to 76 300 adenosine bases for which methylation fraction values were analysed). Since the vast majority of the adenosines present in GATC motifs were fully methylated in all sequenced strains, we analyzed the subset of GATC motifs which exhibited low methylation level in at least one strain as described in Bruneaux et al. 2021: 483 palindromes corresponding to 966 adenosines were used for association analysis with phenotypes (1.2% of all the adenosines in GATC motifs). The significance of the association between each of these 966 epiloci and a given phenotypic trait was calculated as the *p*-value for Spearman’s *ρ* correlation cofficient between the phenotype values and the m6A methylation fractions for the 29 sequenced strains. We used Spearman’s *ρ* (i.e. rank correlation) to avoid excessive leverage from extreme phenotypic values.

To gain insights into potential functional roles of those epiloci, they were associated with annotated genes. A gene was assigned to an epilocus if the adenosine base was located within the gene coding region, or less than 500 base pairs upstream of the initiation codon in order to cover potential regulatory regions of the gene. Several gene set approaches were then tested to try to detect biological functions or pathways related to the epiloci associated with phenotypic traits. We used gene-ontology enrichment tests as implemented in the TopGO R package and KEGG pathway analysis with Wilcoxon rank-sum statistics to compare gene sets, but mostly only very general biological functions were detected with those approaches, such as amino acid or carbon metabolism, nutrient transport and translation (data not shown). Since those approaches are targetting the detection of changes affecting a given biological function or pathway on average, but are not efficient to detect single genes which might affect phenotype, we decided to generate lists of top candidate genes associated with each phenotypic trait (using uncorrected *p*-value*<* 0.005 for Spearman’s *ρ* correlation as the threshold) and to manually curate those genes. Manual curation entailed a literature search to provide a brief description of the function of the gene product in bacterial species and to flag genes potentially involved in chosen categories of interest: regulation of transcription, nutrient transport, excretion, cell wall structure, virulence and a larger last category embracing motility, biofilm formation, adherence and quorum sensing.

## Results

### Identification of a temperature-sensitive prophage in *S. marcescens* genome

In-silico analysis of the genome of the reference strain (ATCC 13880) with PHASTER (Arndt et al., 2016) predicted the existence of five prophage candidates, with two of them incomplete based on PHASTER completeness score (Supplementary Table S1). Using our qPCR-based method to determine phage release rates in liquid cultures under a variety of temperature assays and prophage-specific primers targetting all five prophage candidates, we detected extracellular phage particles only for prophage PP4 across the sequenced clones (Figure 2A). The release of PP4 particles by *S. marcescens* was temperature-sensitive: on average the lowest rates were observed at 38 °C and the highest rates at 24 °C and 31 °C (Figure 2B). From a phage genomics point of view, a comparative analysis showed that PP4 had very high sequence similarity with the *S. marcescens* phage KSP20 previously isolated from aquatic environment (Matsushita et al., 2009) and to P2-like temperate phages and prophages from Enterobacterales (Supplementary Figure S3). We did not find any known virulence factors among the identified proteins of PP4; however, 12 of its 44 predicted proteins remained unannotated (Supplementary Figure S3).

**Figure 2:**
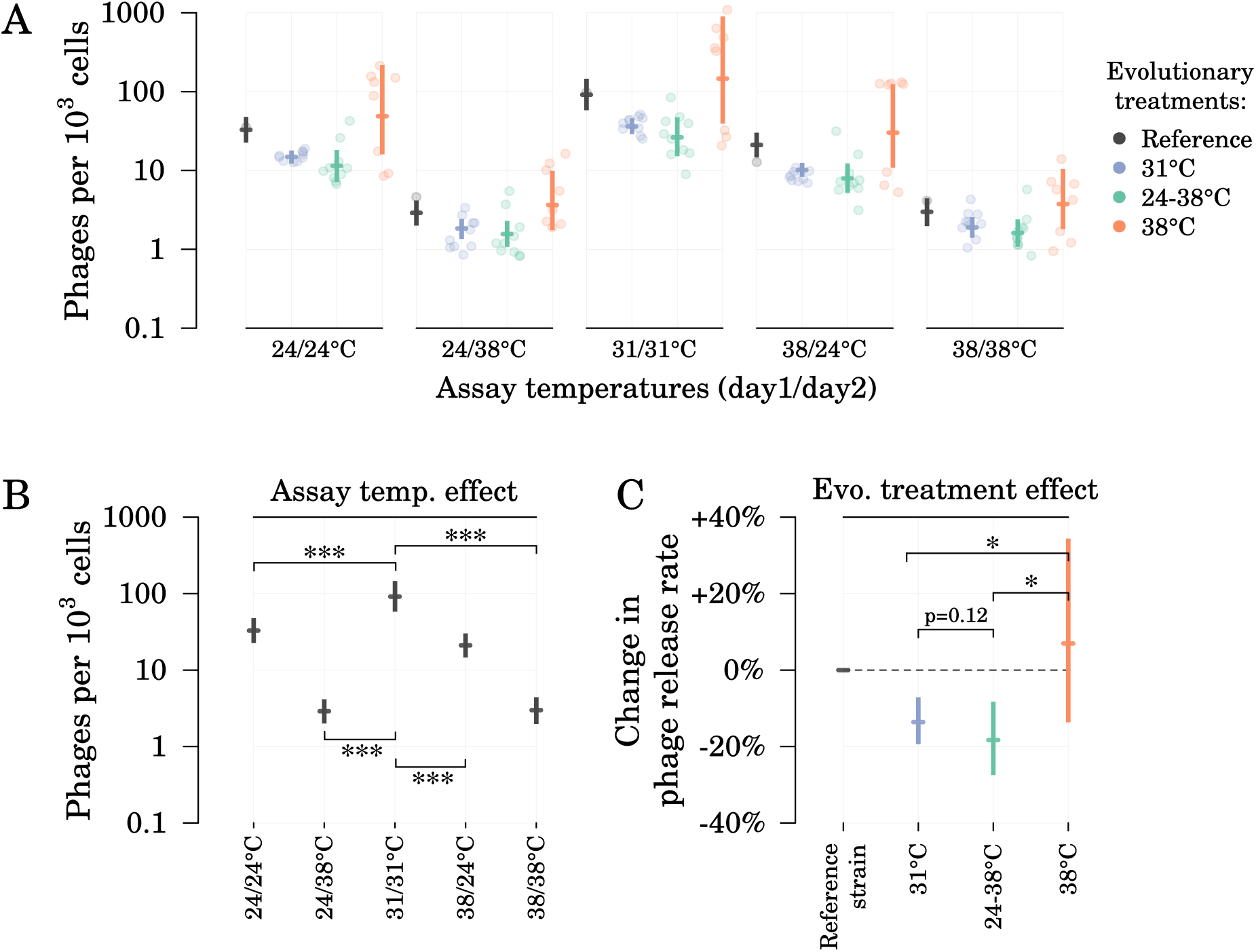
Effect of evolutionary treatment and assay temperatures on the release rates of prophage PP4. Assays lasted two days and assay temperatures are given as day1/day2. (A) Posteriors of the model-estimated mean for each treatment/assay combination. Points are estimated phage release rates for each of the 29 sequenced clones. Full posterior estimates for individual strains are shown in Supplementary Figure S4. (B) Estimates of the assay temperature effects and (C) estimates of the evolutionary treatment effects, with the reference strain used as a reference point. Posteriors are shown as median and 95% credible interval. One-sided Bayesian p-values for pairwise comparisons denoted by * (*p <* 0.05) and *** (*p <* 0.001).

We performed two additional experiments using independent approaches to confirm the presence of PP4 phage particles in our liquid cultures. The first approach was to search for phage particles from pellets prepared from the supernatant of a reference strain culture grown at 31 °C using transmission electron microscopy (TEM). Particles of shape and size compatible with a P2-like phage were successfully observed, albeit in low frequency (Figure 3). The phage identity was confirmed by qPCR using PP4- specific primers on the TEM pellets. The second approach consisted in plaque assays with supernatants from reference strain cultures spread onto lawn cultures of several candidate strains, including the reference strain, but no plaque was observed in any of the conditions tested.

**Figure 3:**
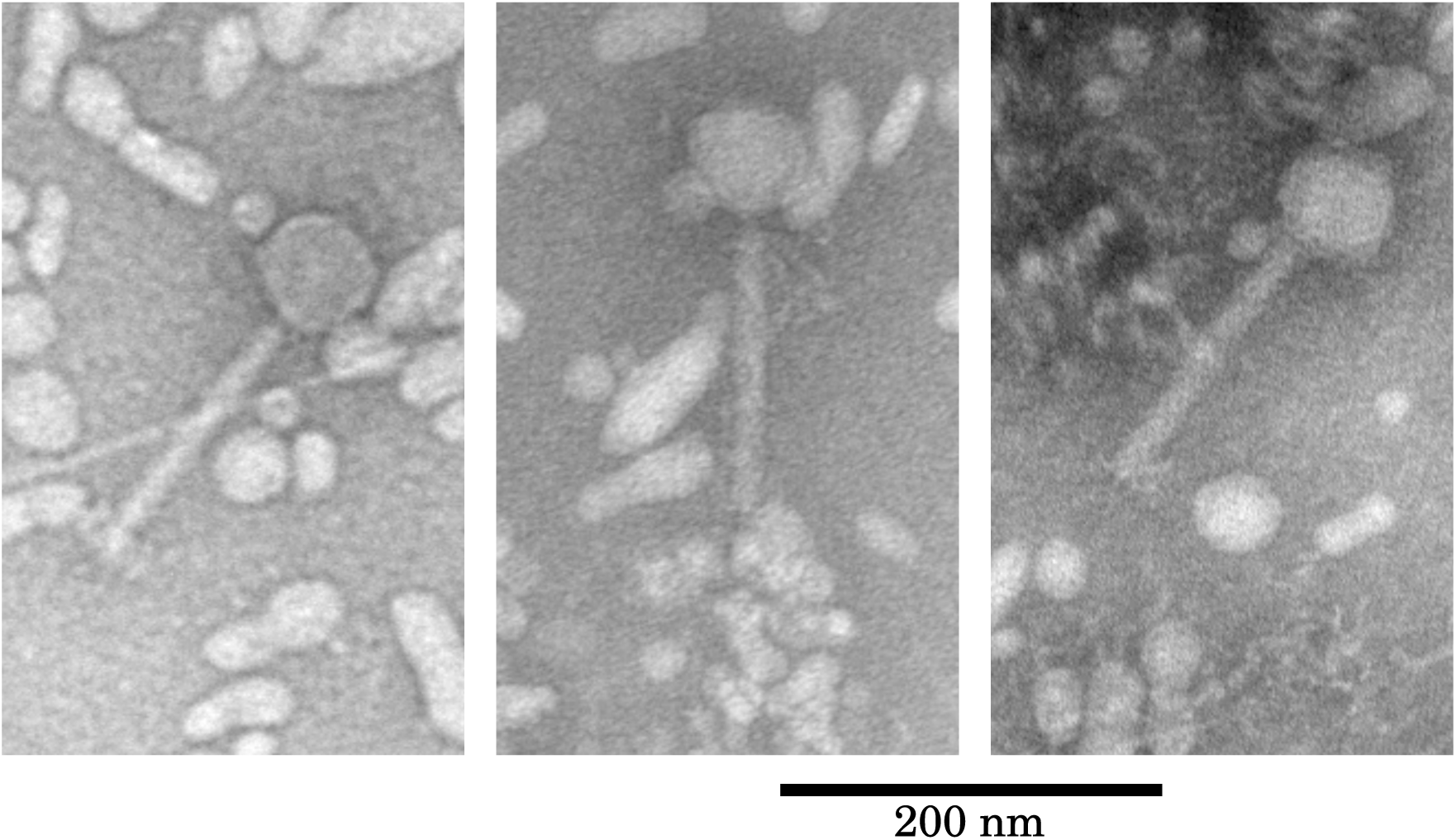
Putative PP4 phage particles observed in transmission electron microscopy (TEM). Negative staining with phosphotungstic acid (PTA) was used. All panels are shown to the same scale.

### Evolutionary changes in phage release rates

By using two-day thermal assays to measure phage release rates (Figure 2A), we evaluated both the effect of mean temperature and of temperature fluctuations on prophage induction in the evolved clones. We found an evolutionary adjustment of prophage release rates in clones evolved under cooler environments (evolutionary treatment effect, Figure 2C). The strains evolved at 31 °C and at 24-38 °C released 19% and 24% less phages, respectively, than strains evolved at 38 °C. Mean patterns of PP4 induction did not differ significantly between strains that had evolved at lower mean temperature (31 °C versus 24-38 °C). The main driver of prophage induction in our assays was the temperature on the final day of the assay, rather than whether a temperature change was experienced during the assay (assay temperatures effect, Figure 2B). Ending an assay at 31 °C induced about three times more phages than ending an assay at 24 °C, and ending an assay at 24 °C induced about ten times more phages than ending an assay at 38 °C.

### Evolutionary changes in bacterium virulence

We estimated the virulence of the experimentally evolved strains by measuring the survival time of waxmoth larvae (*Galleria mellonella*) infected by an injection of 5 μl of bacterial cultures and placed into two assay environments: 24 °C and 31 °C (Supplementary Figure S5). We did not use 38 °C as this temperature would have been lethal to waxmoth larvae. We estimated the relative virulence of sequenced strains from larvae survival data using a Cox proportional hazards mixed model which controlled for both larva body mass and initial density of bacterial cultures and which allowed for different variances across evolutionary treatments (Figure 4A). Those estimates revealed that the average virulence of clones evolved at high temperature (38 °C) tended to be higher than for clones evolved at lower mean temperature when larvae were incubated at 24 °C (38 °C versus 24-38 °C, one-sided Bayesian *p*-value = 0.057; 38 °C versus 31 °C, *p* = 0.066; Figure 4B). However, when larvae were incubated at 31 °C, these differences disappeared while clones evolved at 31 °C were more virulent than those evolved at 24-38 °C (*p <* 0.01). This was consistent with previous results from Ketola et al. (2013) showing that the clones evolved at 31 °C were more virulent than clones evolved at 24-38 °C in a *Drosophila* model. To confirm the results obtained from the 29 sequenced clones, we also measured virulence in waxmoth larvae for a much larger pool of evolved clones from the same original evolution experiment (Figure 1) which confirmed that overall clones evolved at 38 °C had indeed a higher virulence than the others when assayed at room temperature (*p <* 0.01 for comparisons of 38 °C clones with both 24-38 °C and 31 °C clones, Figure 4C). Finally, we observed a moderate to strong positive correlation between average strain virulence in waxmoth larvae and average PP4 release rates (Spearman’s *ρ* = 0.52, *p* = 0.004).

**Figure 4:**
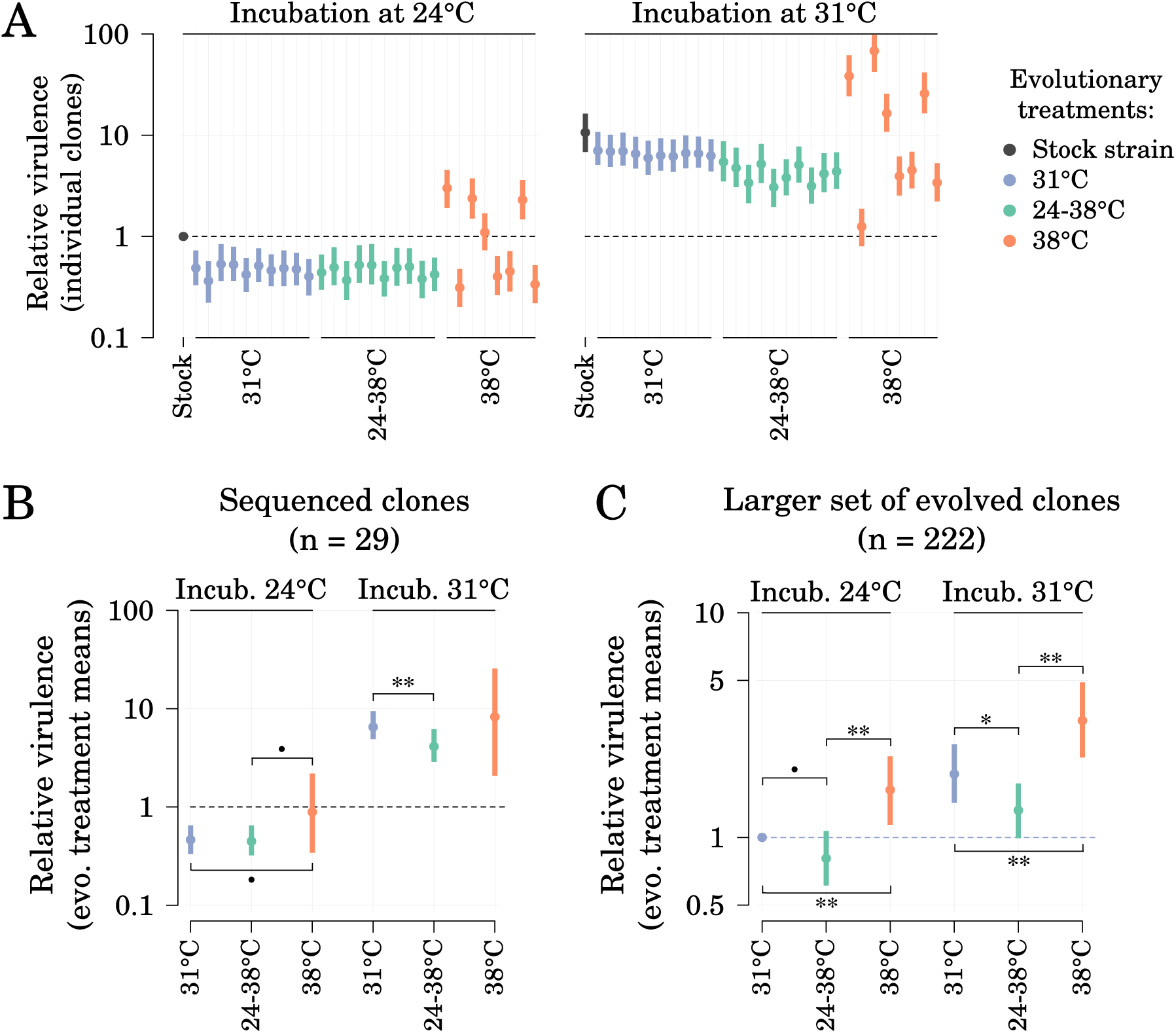
Effect of evolutionary treatment on strains virulence in waxmoth larvae at two incubation temperatures. (A) Relative virulence of individual sequenced clones, measured as relative hazards estimated from a Bayesian implementation of a Cox proportional-hazards model. All virulence estimates are relative to the virulence of the reference strain in incubation at 24 °C (denoted by a broken horizontal line) and are corrected for the effects of injection batch, larval body mass and optical density of injected cultures. (B) Mean relative virulence per evolutionary treatment and per incubation temperature as estimated by the model (exp(*µ_evo_*)) (*n* = 29 sequenced clones). (C) Confirmatory results from a similar virulence experiment utilizing more bacterial clones from the same original evolution experiment (*n* = 222 clones, virulence relative to the average virulence of the clones evolved at 31 °C when incubated at 24 °C). For each model parameter, 95 % credible interval and median of the posterior are shown.

### Association between genetic and epigenetic variation and phenotypic changes

Based on a genome alignment, 52 variable genetic loci were identified among the sequenced clones but none was located inside prophage PP4 sequence (Figure 5 and Supplementary Table S3; Bruneaux et al. 2021). We investigated the association between phenotypic traits and genetic variants present in at least two strains using Wilcoxon rank sum tests adjusted for false-discovery rate (Benjamini and Hochberg, 1995); we also used the methylation data obtained from PacBio SMRT sequencing to test for association between phenotypic traits and adenosine methylation in GATC motifs (which are recognized by an adenine methyltransferase in *S. marcescens*; Ostendorf et al. 1999; Bruneaux et al. 2021). Our main results are described below, and some further details are available in the Supplementary Results section.

**Figure 5:**
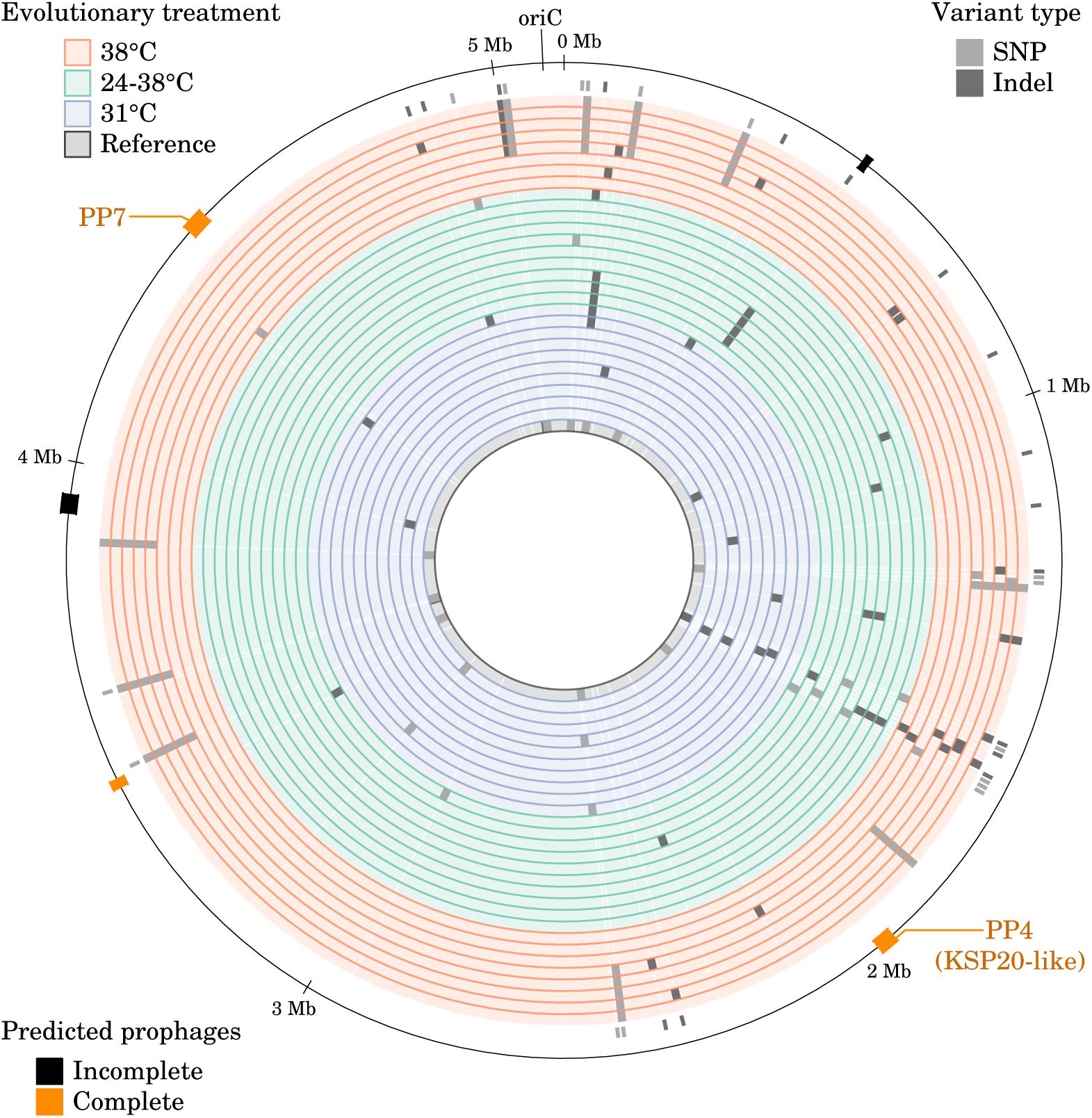
Alignment of the genomes from the 29 sequenced strains showing the variable genetic loci. Each circular track represents a sequenced genome, for which the evolutionary treatment is color-coded. Minor alleles for genetic variants are shown on the genome tracks in light grey (SNP) and dark grey (indels). Ticks outside the last genome track indicate non-synonymous variants (i.e. non-synonymous SNPs and in-dels resulting in a frame shift). The outer line represents coordinates along the genome and the locations of the five predicted prophages. Prophages PP4 and PP7 are the prophages shown in Figure S3.

Several of the variable genetic loci associated with prophage induction and with bacterial virulence had a potential role in the biofilm structure and in the outer structure of the cellular envelope in (e.g. genes involved in peptidoglycan and LPS biosynthesis; Figure 6 and Supplementary Table S3). A striking pattern was the presence of three distinct mutations occurring in a single glycosyltransferase gene and close to the putative active site of the protein (mutations *28*, *29* and *30*, Supplementary Figure S6, Supplementary Table S3). These mutations were observed independently in three strains evolved at 24-38 °C and in one strain evolved at 38 °C (which had low prophage induction rates and low virulence compared to the other strains evolved at 38 °C).

**Figure 6:**
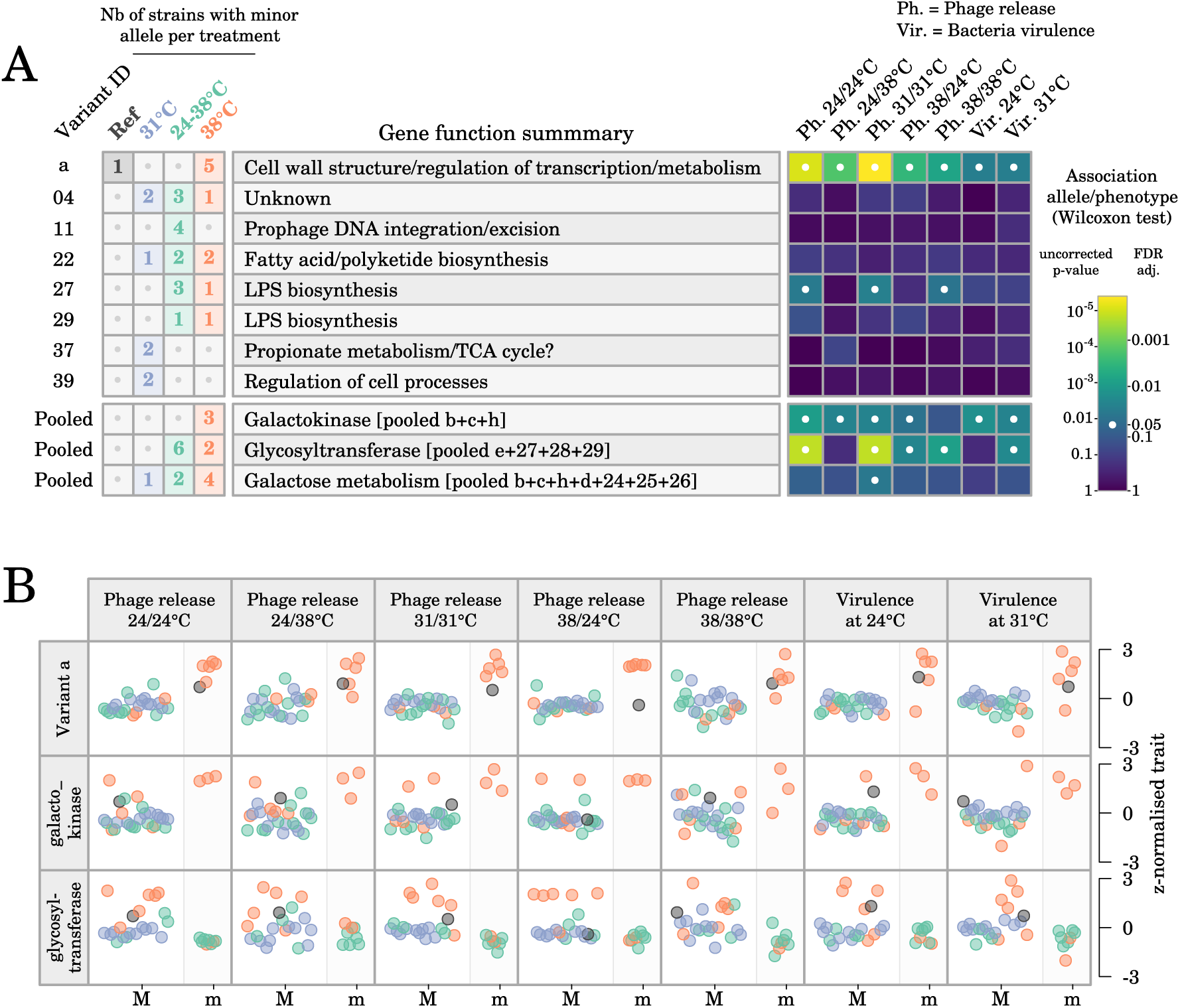
Association between phenotypes and genetic variants observed in at least two sequenced strains. (A) Distribution of genetic variants across evolutionary treatments and association between alleles and phenotypes based on Wilcoxon rank sum tests. Variant IDs can be matched with those in Supplementary Table S3 for details. (B) Visualization of the association between phenotypic values and major (M) and minor (m) alleles of variant *a* and of the “pooled” variants for galactokinase and glycosyltransferase. Colors correspond to the evolutionary treatment applied to each strain.

Fity-two genes were found associated with phenotypic changes via adenosine methylation changes and we manually curated their functional annotation based on literature (Figure 7). Many of those genes were involved in functional categories that are critical for pathogen virulence in other bacteria species, such as nutrient capture, excretion into the outer medium, biofilm formation, adherence, and motility, all of which have a key role in successful colonization and invasion of the host tissues (Turner et al., 2009; Luo et al., 2017; Liu et al., 2017; Ren et al., 2018) (Figure 7). Additionally, several genes were involved in cell wall structure, notably in lipopolysaccharide biosynthesis.

**Figure 7:**
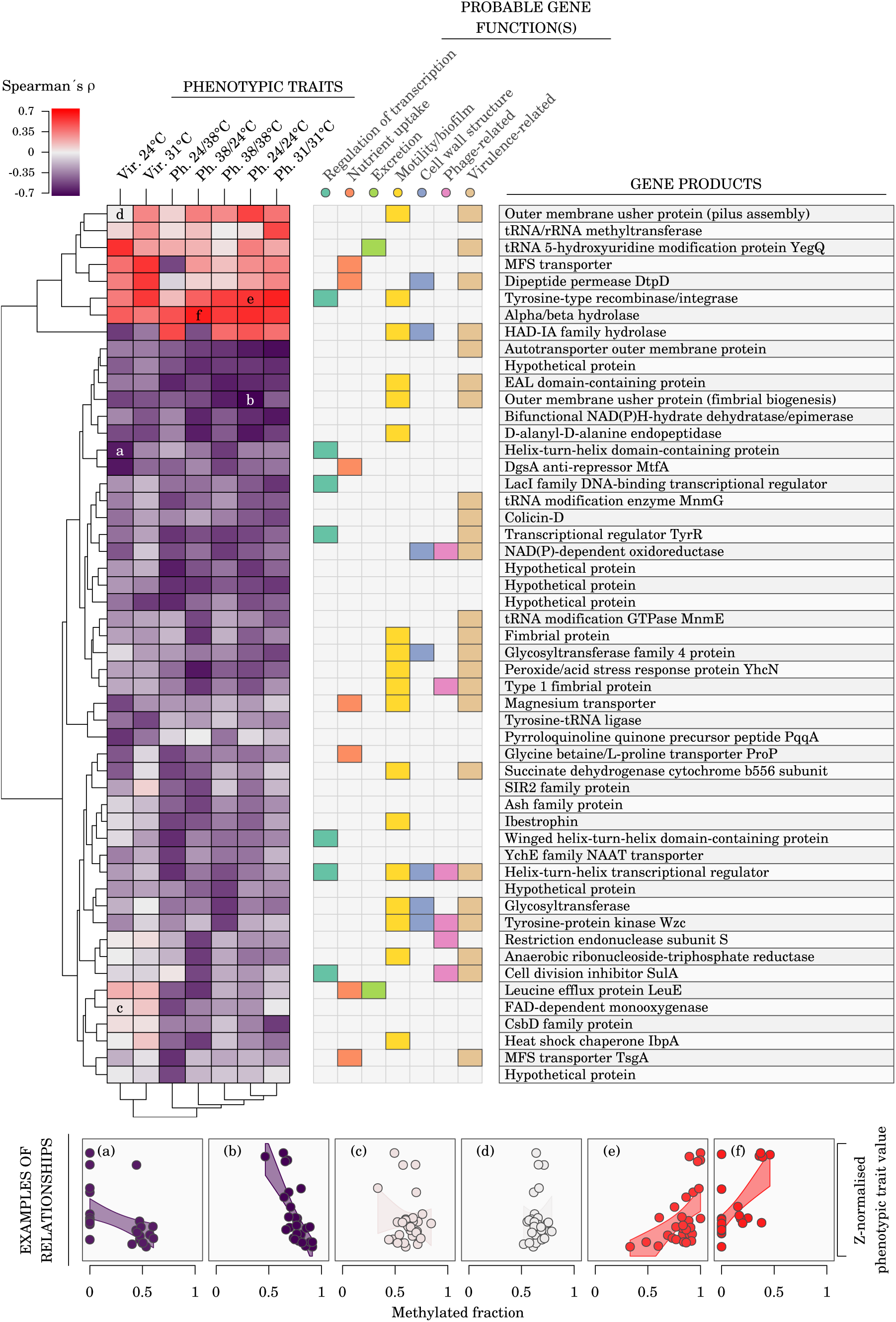
Association between phenotypic traits and adenine methylation changes. The heatmap shows Spearman’s *ρ* between methylated fractions of variable m6A epiloci (rows) and phenotypes (columns; Ph., phage release; Vir., bacteria virulence in waxmoth larvae). Overlapping or closest (*≤* 500 bp) down-stream genes were assigned to each m6A epiloci. Probable gene functions were assigned to each gene product based on a manual literature search. Visualization of the relationship between m6A methylated fractions and phenotypic trait values is shown for the heatmap cells marked with a letter. The m6A epiloci used here were the variable m6A epiloci with a methylation fraction range *≥* 0.2 across sequenced samples and an uncorrected *p*-value *≤* 0.005 for Spearman’s *ρ* with at least one phenotypic trait.

## Discussion

### *S. marcescens* carries a P2-like, temperature-inducible prophage

We used three independent methods (qPCR, TEM, plaque assays) to establish if any of *S. marcescens* prophages could result in phage particles in our experiments. Taken together, our results confirmed that the PP4 prophage is induced in *S. marcescens*cells and released as extracellular phage particles at a low, temperature-sensitive rate as detected by qPCR and confirmed by TEM, but suggest that the burst and/or infectivity rates of this phage are low enough that no clearing is observed in our plaque assays. Interestingly, the immunity repressor protein located immediately upstream of the PP4 integrase is of the short type (99 amino-acid long) and thus most likely of the P2-type rather than of the longer 186-type (Christie and Calendar, 2016). Since phage 186 is inducible by the SOS response while phage P2 is not, this suggests that prophage PP4 might be non-inducible by the SOS response like P2 (Christie and Calendar, 2016; Lamont et al., 1989).

Structural proteins were well conserved between PP4 and related sequences, as is common for P2-like phages (Nilsson and Haggåard-Ljungquist, 2007), with the exception of tail fiber proteins (Figure S3). Tail fiber proteins are known to be involved in phage host specificity (Scholl et al., 2001; Le et al., 2013; Yehl et al., 2019) and P2- like phages, while exhibiting strikingly similar structural genes even when infecting different species, have been suggested to coevolve with their bacterial hosts (Nilsson and Haggåard-Ljungquist, 2007).

### Abiotic changes affects the evolution of prophage-bacterium interaction

Given that *S. marcescens* carries a temperature-inducible prophage, a comparison of phage release rates after bacterial evolution under different temperatures provides a compelling opportunity to explore how environmental changes can affect the evolution of prophage-bacterium interactions. In particular, increased phage release rates are likely to have a negative effect on bacterial fitness in a monospecies population, assuming that increased release rates are at least partly due to increased induction and/or burst rates. It is thus reasonable to expect that selection would favor counter-adaptations in *S. marcescens* to reduce the fitness loss due to prophage induction and cell lysis in cooler environments (Canchaya et al., 2003), while evolution in warmer environments would release selective pressure against such fitness loss. This prediction was confirmed based on the phage release rates we measured for evolved clones: the evolution of lower phage release rates in clones from evolutionary treatments where prophage-inducing temperatures occurred (i.e. treatments 31 °C and 24-38 °C) was consistent with a temperature-dependent effect of PP4 on *S. marcescens* fitness.

### Induced evolutionary changes have consequences for bacterium virulence

After confirming that evolutionary changes occurred in the interaction between prophage PP4 and its bacterial host, we explored how these environmentally-triggered changes might cascade to another biological level: an insect host of the bacterium itself. Previous evolution experiments with the same bacterial species have shown that evolution in the presence of a protist predator or of a lytic phage affected *S. marcescens* virulence in an insect host (Friman et al., 2011; Mikonranta et al., 2012). In particular a trade-off between resistance to a lytic phage and bacterial virulence traits was demonstrated by Friman et al. (2011): the presence of the lytic phage prevented the evolution of more virulent *S. marcescens* clones at 37 °C while virulence of bacteria evolving at the same temperature but without the lytic phage increased. Those studies support the coincidental evolution hypothesis, which suggests that selection pressures acting on an opportunistic pathogen in the external environment can affect the pathogen virulence in its host (Levin and Edén, 1990; Brown et al., 2012). In our experiment, we hypothesised that evolutionary changes related to phage release rates might also impact bacterial virulence.

We observed substantial correlation between average strain virulence in waxmoth larvae and average PP4 release rates, which confirmed that prophage induction in liquid medium closely related to the bacteria virulence in the insect host and is in line with the coincidental evolution hypothesis. Interestingly, the original selection pressure in our case was abiotic (environmental temperature acting on phage release rates), contrary to the protist predator and the lytic phage used by Friman et al. (2011) and Mikonranta et al. (2012).

### Both genetic and epigenetic variation can play a role in phenotypic changes

To explore the mechanistic basis of the evolutionary changes in prophage release rates and in bacterial virulence, and the potential connection between those two traits, we analysed the genomic and methylation data of the 28 evolved strains used in our study and of the reference strain pre-existing the evolution experiment (Figure 1; Bruneaux et al. 2021). We found no known virulence factor among the annotated proteins of PP4, but some accessory proteins found in P2-related phages are known to contribute to bacterial virulence, protection against other bacteriophages, or SOS prophage induction (Christie and Calendar, 2016). More generally, prophage-encoded virulence factors are considered one of the benefits brought by prophages to their bacterial host explaining the maintenance of prophages in bacterial genomes (Fortier and Sekulovic, 2013; Koskella and Brockhurst, 2014). Another possible direct connection between prophage induction and bacterial virulence is via the release of endotoxins upon cell lysis that could be a causative agent of *S. marcescens* virulence since its lysates are known to be cytotoxic on their own (Petersen and Tisa, 2012).

Given that none of the variable genetic loci we observed in our dataset were located inside the prophage PP4 sequence, it can be reasonably expected that mutations elsewhere in the genome and/or epigenetic modifications should be responsible for evolutionary differences in prophage induction rates and bacteria virulence among the sequenced clones. The variable genetic loci we found associated with phage release rates and with bacterial virulence suggest that biofilm formation and outer cell wall structure can have a role in modulating those traits. In particular, the three distinct independent mutations in the glycosyltransferase gene we observed strongly support the hypothesis of a role of the outer cellular envelope in the evolutionary response against phages, with potential consequences on bacterial virulence. Similarly, among the genes for which variation in adenosine methylation was associated with changes in phage release rates or in bacterial virulence, several were related to lipopolysaccharide biosynthesis. All in all, these results indicates that the O antigen, which is typically involved both in cell recognition by phages and in bacteria virulence in their host (Chart et al., 1989; Li and Wang, 2012), could act as an important player in evolutionary trade-offs between bacterial virulence and resistance to phage infection. It has been noted previously that *S. marcescens* strains exhibit a large diversity of O antigens (Gaston and Pitt, 1989), and one particular study showed that *S. marcescens* cells grown at different temperatures had different LPS structure and phage affinity, with cells grown at 37 °C having shorter O antigen and lower affinity for LPS-specific phages than cells grown at 30 °C (Poole and Braun, 1988). Those suggested links are consistent with earlier research: Flyg et al. (1980) showed that phage-resistant *S. marcescens* mutants also had reduced virulence in *Drosophila* and Cota et al. (2015) demonstrated that an epigenetic mechanism altering the O-antigen chain length in *Salmonella enterica* created a trade-off between bacterial virulence and phage resistance.

Finally, a striking feature of the genetic and epigenetic changes observed in our study was that almost none of the associated genes was directly related with thermal selection pressure (only one heat shock chaperone was found, *IbpA*; Figure 7), even though temperature was the primary selective pressure in the evolution experiment. For instance, we did not find indication of changes related to other HSP or to DNAK genes, even though they are known to be the target of selection in hot and fluctuating environments (Sørensen et al., 2003; Ketola et al., 2004; Sørensen et al., 2016). Such a weak direct effect of the experimentally manipulated factor underlines the fact that indirect selection due to prophage induction is likely to have overruled the direct effects of temperature. It is also consistent with the fact that the temperature treatment during evolution was not strongly asssociated with genetic or epigenetic changes (Bruneaux et al., 2021), but more associations are observed when we compare directly genetic and epigenetic data and phenotypic traits (which allows to capture the phenotypic variation between clones within an evolutionary treatment).

## Conclusion

We showed that the opportunistic pathogen *S. marcescens* harbored a temperature-sensitive prophage and that phage release rates were susceptible to evolutionary changes. Consistent with the hypothesis that phage release decreased bacterial fitness (at least in laboratory conditions), we observed compensatory evolutionary trajectories where intrinsic release rates decreased in bacteria evolving under strongly release-inducing temperatures (31 °C and 24-38 °C) while intrinsic release rates increased in some bacteria evolving under weakly release-inducing temperature (38 °C). Our experiments did not allow us to clearly identify which phage properties were affected by culture temperature (induction rate, burst size, capsid stability) and which molecular mechanisms were responsible for the evolution of different phage release rates in *S. marcescens*, but the genetic and epigenetic data were compatible with an important role of the structure of the outer cell wall. We also observed evolutionary changes in *S. marcescens* virulence, which were correlated with the changes in intrinsic phage release rates. Again, our genetic and epigenetic data are suggestive of mechanistic links probably involving outer cell wall structure, even if more research is needed to confirm the actual molecular mechanisms involved.

Our results provide an original instance of coincidental evolution where an abiotic environmental factor driving (pro)phage release rates in a bacterium had cascading effects on the evolution of bacterial virulence. Such occurrences of coincidental evolution can be hard to predict but are important to take into account to improve our understanding of the complex feedback loops between environment, viruses, microbes and ecosystems.

## Acknowledgements

We acknowledge Kati Saarinen and Lauri Mikonranta for help with the virulence experiments, Lotta-Riina Sundberg and Leena Lindström for comments on the manuscript, the Academy of Finland (Project 278751) and the Centre of Excellence in Biological Interactions for funding and facilities and the CSC – IT Center for Science, Finland, for computational resources used in this project.

## Data accessibility and benefit-sharing

### Data accessibility

PacBio sequencing data (HDF5 files) were submitted to the European Nucleotide Archive’s Sequence Read Archive (ENA-SRA, https://www.ebi.ac.uk/ena, project PRJEB40306) and assembled genomes were submitted to NCBI’s GenBank (biosamples SAMEA7301478 to SAMEA7301506). Genetic, epigenetic and phenotypic datasets used for analysis will be submitted to the Dryad repository (https://datadryad.org/stash).

Availability of biological materials: The evolved clones of *Serratia marcescens* used in this study are available from the authors upon request.

### Benefits generated

Benefits from this research accrue from the sharing of our data and results on public databases as described above.

## Authors contributions

TK, JAG and IK conceptualized the study. Experimental design for PacBio sequencing was done by TK and MB. DNA extraction for PacBio sequencing was done by RA. AMÖO identified prophage sequences. AMÖO, RA and MB designed the prophage induction assays and RA and MB performed the assay experiments. EL performed the electron microscopy experiments and the plaque assays and helped analyze PP4 sequence. ZC performed virulence experiments of the sequenced clones in the insect host. RA, IK, MB, and TK performed virulence experiments for the larger pool of evolved clones. MKS assisted with laboratory experiments. MB and IK analysed the sequencing data. MB, IK and TK wrote the original draft with later edits and reviews by all co-authors.

## 1 Supplementary Results

### 1.1 Association between genetic changes and phenotypic traits

The variable loci most clearly associated with both phage release and bacteria virulence in the insect host were the variant *a* and the pooled variants related to galactokinase (pooled variants *b*, *c*, and *h*) and related to glycosyltransferase (pooled variants *e*, *27*, *28*, and *29*) (Figure 6). These genetic variants were located in or close to (*≤* 500 bp) genes annotated as transcriptional regulators (molybdenum-dependent transcriptional regulator and transcriptional regulator RcsB involved in motility and capsule and biofilm formation in *E. coli*) and enzymes involved in the cell wall and outer membrane structure and biofilm formation (peptidoglycan synthase, two glycosyltransferases and a cellulose biosynthesis protein BcsG) (Supplementary Table S3). Those genes point towards a potential role for modifications of biofilm structure and of the outer structure of the cellular envelope in modulating phage particle production and virulence in the insect host. In particular, the three independent mutations located in a single glycosyltransferase gene (mutations *28*, *29* and *30*, close to the putative active site of the protein; Supplementary Figure S6; Supplementary Table S3) were observed independently in several strains: three strains evolved at 24-38 °C and one strain evolved at 38 °C. Those independent mutations point to the important role of the outer cellular envelope in the evolution against phage.

Finally, we also noted that haplotype *a*, comprising eleven associated genetic loci, was shared by 5 out of the 8 strains evolved at 38 °C and by the reference strain, but by none of the other sequenced strains. This points to the probable existence of some standing genetic variation at the onset of the experiment, which was then subjected to selection during the experimental evolution (Bruneaux et al., 2021).

### 1.2 Association between epigenetic changes and phenotypic traits

In addition to nucleotide sequences, the data we obtained from the PacBio SMRT method also provided information about base methylation. In *S. marcescens*, adenosines present in GATC motifs are methylated into m6A by the Dam enzyme at a very high rate (*>*98% of GATC motifs were methylated on both strands in our dataset; Bruneaux et al. 2021). The remaining GATC motifs can be either hemi-methylated or unmethylated in a cell, and are often variably methylated across cells of a given culture and across strains. Adenosine methylation can influence gene expression by affecting the binding of regulatory proteins to promoter regions of genes (Gomez-Gonzalez et al., 2019) or by affecting transcription speed via increased DNA stability of gene bodies (Riva et al., 2004a,b). Such epigenetic regulation can be maintained across rounds of DNA replication by competitive binding to target DNA between the Dam responsible for methylation and regulatory proteins specific to the same region (Casadesús and Low, 2006, 2013), and can thus be subject to selection.

Among GATC motifs which were not fully methylated in our dataset, no association was found between evolutionary treatments and methylated fractions (Bruneaux et al., 2021). However, we identified adenosines for which changes in methylation level were associated with phenotypic changes in the traits measured here (phage induction and virulence in an insect host). For a given phenotypic trait, GATC loci exhibiting both positive and negative correlations between methylated fractions and the trait values could be observed (Figure 7, heatmap panel). Manual curation of the function of the genes associated with GATC motifs related to phenotypic changes showed that many of them were involved in (1) transcription regulation, (2) nutrient capture and transport into the cell, (3) excretion into the outer medium, (5) biofilm formation, adherence or motility, and (6) cell envelope structure (including peptidoglycan and lipopolysaccharide biosynthesis) (Figure 7, gene functions panel). Many of those functional categories have been shown to be critical for pathogen virulence in other bacterial species, in particular for nutrient capture in the challenging host medium (Ren et al., 2018; Liu et al., 2017), for recognition of the host habitat via its nutrient signature (López-Garrido et al., 2015; Krypotou et al., 2019) and for biofilm formation, adherence and motility which have a key role in colonization and successful invasion of the host tissues (Turner et al., 2009; Luo et al., 2017). The numerous candidate genes involved in lipopolysaccharide biosynthesis also suggest that the O antigen, which can classically be involved both in cell recognition by phages and in bacterial virulence in its host (Chart et al., 1989; Li and Wang, 2012), could act as a major player of evolutionary trade-offs between bacterial virulence and resistance to phage infection.

## 2 Supplementary Methods

### 2.1 Quantification of phage induction using qPCR

#### 2.1.1 Culture conditions for the temperature assays

Frozen stocks had been stored at *−*80 °C in 40 % glycerol, with evolved clones stored in 100-well plates (Bioscreen measurement plates), in randomized order and reference clone stored in microcentrifuge tubes. A preculture step in 400 μl of SPL 1 % at 31 °C was performed by using a cryo-replicator to inoculate evolved clones into a new 100- well plate and by inoculating the reference strain into wells of another plate. After 24 hours, five identical 100-well assay plates containing both the 28 evolved clones of interest and the reference clone were prepared by transferring 40 μl of each preculture into 360 μl of fresh SPL 1 % (1 well per clone, i.e. 29 wells occupied per plate). For the first day of assay, one plate was incubated at 31 °C, two plates at 24 °C and two plates at 38 °C. After 24 hours, clones within a given plate were transferred to 29 previously empty new wells in the same plate (40 μl culture into 360 μl fresh medium). For the second day of assay, the plate from 31 °C was kept at 31 °C, one plate from 24 °C was kept at 24 °C and the other was transferred to 38 °C, and one plate from 38 °C was kept at 38 °C while the other was transferred to 38 °C. After 24 hours, plates were taken for sample processing. Extra wells containing sterile SPL 1 % medium were used on the assay plates to monitor potential contamination during plate handling (which was not observed). The whole experiment was performed twice, starting with the same frozen stocks but with independent precultures.

#### 2.1.2 Sample processing and qPCR runs

At the end of the second day of assay, each of the 29 cultures in each of the 5 assay plates was processed in the following way: 50 μl of native culture sample was transferred to a 96-well PCR plate, while the rest of the culture was placed into a microcentrifuge tube, centrifuged at 13 500 g for 5 min and 50 μl of supernatant was transferred in the 96-well plate, resulting in two paired samples per culture (native and supernatant). Samples from a given assay plate were placed into the same 96-well plate. A DNase treatment was then performed to digest DNA fragments which were not protected inside a bacteria cell or a phage particle. 5 μl of DNase I at 0.1 mg ml*^−^*^1^ were added to each sample, followed by an incubation at 37 °C for 30 min. DNA was then released from bacteria cells and potential phage particles by incubating the samples at 95 °C for 30 min after having added 5 μl of EGTA (20 mm, pH 8) in order to hinder DNase I activity. The sample plates were then stored at *−*20 °C until DNA quantification by qPCR runs.

Quantification of DNA target sequences was performed using prophage-specific primer pairs and one bacterial-gene-specific primer pair (Supplementary Table S2). Preliminary experiments using the reference strain at 31 °C having showed no detectable extra-cellular DNA at least for prophages 2 and 5, six qPCR were runs per 96-well sample plate from this experiment using primers for prophages 1, 3, 4, 6, 7 and for bacterial gene purA2. Runs were performed using CFX Real Time PCR detection system (Bio-Rad laboratories, USA). Amplifications were performed in a final volume of 10 μl, containing 5 μl of 2 x IQ SYBR Green Supermix (Bio-Rad), 0.5 μl of forward and reverse primers (6 μm each) and 1 μl of undiluted sample. Amplifications for each primer pair were performed on separate qPCR plates, with in-plate calibration samples for each run. Calibration samples were prepared by serial dilution of a stock solution of purified *Serratia marcescens* DNA of known concentration, and ranged in concentration from 10^6^ to 1 genome copy per qPCR well, based on the predicted molecular weight of *S. marcescens* chromosome. Experimental and calibration samples were run in triplicates within each qPCR plate. The qPCR reaction used an initial denaturation step lasting 3 min at 95 °C, followed by 41 cycles consisting of denaturation at 95 °C for 10 s, annealing at 61 °C (for all prophage primers) or 56 °C (for bacterial gene primers) for 10 s, and elongation at 72 °C for 30 s. A melt curve analysis was performed at the end of the run to check the quality of the amplified product (from 65 °C to 95 °C, using increments of 0.5 °C and 5 s steps). In-plate calibration samples were used to estimate the efficiency *E* of the qPCR reaction with the formula *E* = *−*1 + 10^(^*^−^*^1^*^/β^*^)^ where *β* is the slope of the linear relationship between *Cq* values and log_10_(concentration) for the calibration samples. To test for an effect of potentially undegraded RNA molecules on phage activation estimates, some samples were incubated with RNase for 30min just prior to qPCR runs. Estimates of phage activation for those samples were similar whether the samples were treated or untreated with RNase prior to qPCR runs, hence data from both RNase-treated and untreated qPCR runs was used for downstream analysis.

#### 2.1.3 Estimation of prophage induction rates and treatment effects using a Bayesian model

We incorporated into a single Bayesian model the simultaneous estimation of phage induction rates and of the effects of assay temperature and evolutionary treatment. To simplify its presentation here, we will first explain the modelling part related to the estimation of induction rates for each culture well, based on the Cq values for the native and supernatant samples obtained from qPCR runs with bacterial and prophage primers, before explaining the incorporation of assay and evolutionary treatment effects.

Let *c_bact,nat_* be the number of bacterial chromosome copies present in a native sample. The value of *c_bact,nat_* is determined from the qPCR run using the bacterial-gene-specific purA2 primers. Let *c_pro,nat_* be the number of prophage DNA copies present in the native sample for e.g. prophage KSP20. The value of *c_pro,nat_* is determined from the qPCR run using the prophage-specific primers. Let’s assume that this prophage is induced into phage particles at an activation rate *a*, such that the number of phage particles present in the native sample *c_phg,nat_* is related to the number of bacteria cells (i.e. the number of bacteria chromosome copies) by *c_phg,nat_* = *a × c_bact,nat_*. Since the prophage primers can target the prophage sequence both in the bacterial genome and in phage particles, we have:

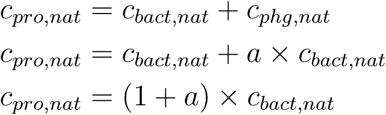

After centrifugation, we assume most bacteria cells have been pelleted and most phage particles (if any) have remained in suspension. Let *k* be the concentration factor during centrifugation for this culture, so that *k* = (*c_bact,sup_/c_bact,nat_*) where *c_bact,sup_* is the number of bacterial chromosome copies present in the supernatant samples, as determined by qPCR with purA2 primers (0 *≤ k ≤* 1). If *c_pro,sup_* is the number of prophage DNA copies in the supernatant sample determined by qPCR with the prophage primers and *c_phg,sup_* is the number of phage particles in the supernatant sample, and if we assume *c_phg,sup_* = *c_phg,nat_* (i.e. we assume that the amount of phage particles pelleted during centrifugation is negligible), we have:

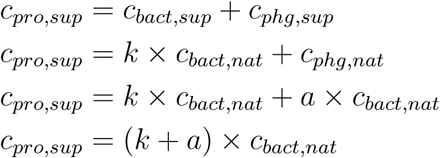

Thus, to summarize, the two fundamental equations that relate the four qPCR measurements for a given culture (*c_bact,nat_* / *c_bact,sup_* / *c_pro,nat_* / *c_pro,sup_*) and the prophage activation rate *a* in this culture are:

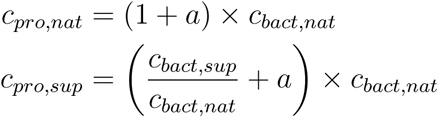

We describe below the integrated Bayesian model used to estimate phage activation rates based on those two equations and on the Cq data obtained from qPCR runs. Note that in the model description below, all parameters corresponding to DNA concentrations are expressed in number of target copies per qPCR well (copies/well).

The model relates Cq values to DNA concentrations, using plate-specific calibration parameters (calibration samples were present in all qPCR plates). Firstly, the model likelihood component due to the calibration samples is (with *n_cal_* being the total number of qPCR wells containing a calibration sample in our dataset):

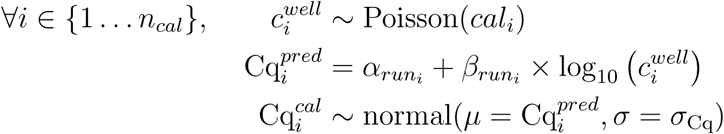

where *cal_i_* is the expected number of target copies in the well (between 1 and 10^5^ in our experiment), 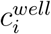 is the actual number of target copies in the well, *α*_[_*_·_*_]_ and *β*[.] are calibration parameters describing the relationship between Cq values and DNA concentrations, *run_i_ ∈ {*1 *… n_runs_}* is the index of the qPCR plate corresponding to the calibration sample and 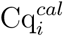 is the observed Cq value for the sample. Note that *α*_[_*_·_*_]_ and *β*_[_*_·_*_]_ are plate-specific (i.e. they are indexed by *run_i_*) to account for plate variability in the qPCR efficiency, while the parameter *σ*_Cq_ which accounts for the experimental noise in the observed Cq values is shared across all qPCR runs. Note also that we use a Poisson distribution for 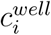 to more accurately describe the sampling process happening when pipetting the template from their preparative tubes into the qPCR wells, especially at low template concentrations.

Secondly, we describe the model likelihood component due to the qPCR wells containing the experimental samples of unknown concentrations prepared from the cultures in the assay plates. For this, we set (with *n_unkn_* being the number of qPCR wells with samples of unknown concentration and *cult_i_* the index of the original culture for each unknown sample):

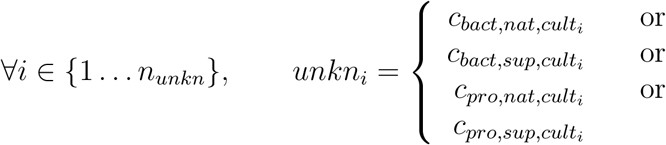

depending on whether the unknown sample is run with purA2 (*c_bact,.,._*) or prophage (*c_pro,.,._*) primers and whether it is native (*c_.,nat,._*) or from supernatant (*c_.,sup,._*). The likelihood due to unknown samples is then of the same form as for the calibration samples:

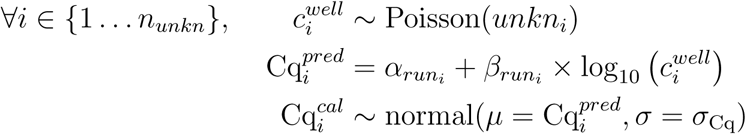

The remaining deterministic relationships of the model and the priors used for unknown parameters to estimate are:

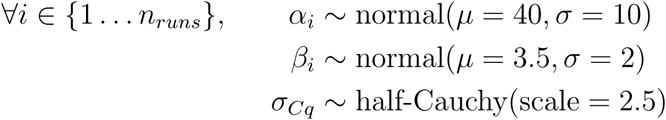

for the parameters of the qPCR calibration curve for each run (note that *σ_Cq_* is shared across all qPCR runs) and:

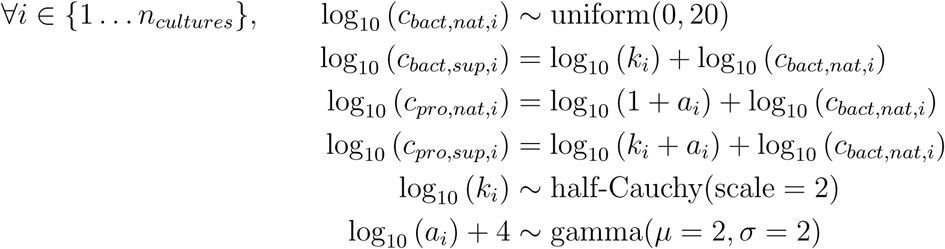

for the characteristics of a given culture well. Note that here, we assume that the minimum value of activation rate *a* is 10*^−^*^4^, which is approximatively the lower sensitivity threshold predicted for our method when we assume that Cq values are measured with a standard deviation *σ_Cq_ ≈* 0.48 (Supplementary Figure S7). We model this as (log_10_ (*a_i_*) + 4) following a Gamma distribution. In this explanation, we use fixed values for the parameters of the Gamma distribution, but when we will introduce the effect of assay and evolutionary treatment the *µ* and *σ* parameters of this Gamma distribution will depend on the treatments.

This model formulation is sufficient to obtain posterior distributions for log_10_(*a_i_*) for each culture well *i* in the assay plates. To model the effect of assay and evolutionary treatment, we extend the model by modifying the parameters of the previous prior for *a_i_*:

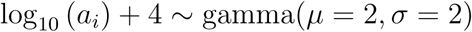

by:

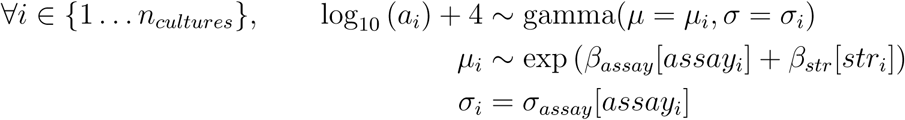

where *assay_i_* is the index of the assay treatment for culture *i* (*assay_i_ ∈ {*1 *…* 5*}*) and *str_i_* is the index of the strain ID for culture *i* (*str_i_ ∈ {*1 *…* 29*}*). The priors for the effect of assay treatments are:

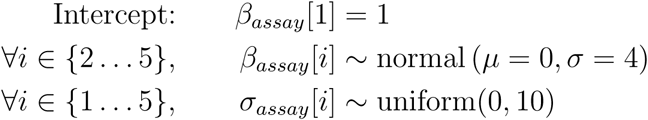

The strain effects include a hierarchical effect of the evolutionary treatment (four levels: three evolution environments plus the reference strain). The priors for the strain and evolutionary treatment effects are:

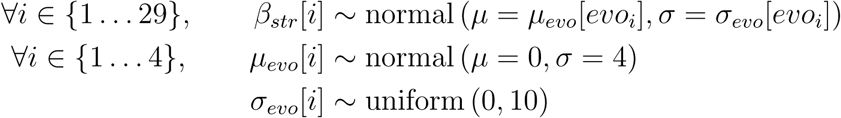

where *evo_i_* is the index of the evolutionary treatment for strain *i*.

### 2.2 Bayesian implementation of the Cox proportional hazards mixed model

The virulence experiment dataset contained observations for *N* = 2182 individual larvae. For each larvae *i*, survival time *s_i_* was calculated as the difference between recorded death time and injection time. The survival timeline for all larvae was divided into *T* = 20 intervals, so that the *s_i,i∈{_*_1_*_…N}_* values were homogeneously distributed across intervals (i.e. all intervals contained approximatively the same number of death events). Intervals were defined by their boundaries *t_j,j∈{_*_1_*_…T_*_+1_*_}_*, such that interval *j* is [*t_j_, t_j_*_+1_) and is of duration *dt_j_*= *t_j_*_+1_*− t_j_*.

The survival data was transformed into a risk variable *Y_i_*(*j*) and an event count variable *dN_i_*(*j*) defined for all *i ∈ {*1 *… N}* and *j ∈ {*1 *… T}* by:

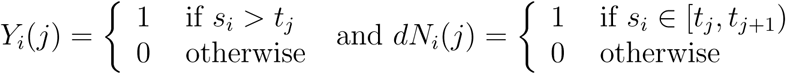

The model assumes:

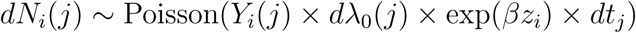

where *dλ*_0_(*j*) is the increment in the integrated baseline hazard from *t_j_* to *t_j_*_+1_ and *βz_i_*is the product of the model parameters and of the covariate values for larva *i*. The term *βz_i_*corresponds to:

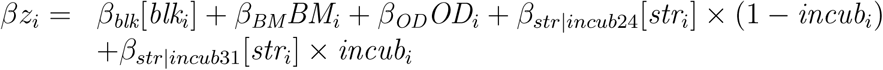

where *blk_i_*, *BM_i_*, *OD_i_*, *str_i_*, and *incub_i_* are respectively the replication block, body mass, preculture OD, injected strain ID (*str_i_ ∈ {*1 *…* 29*}*) and incubation temperature (0 for 24 °C, 1 for 31 °C) for larva *i*. Square brackets indicate indexing of a vector parameter; *β_blk_*is a vector containing the replication block effects and *β_str|incub_*_24_and *β_str|incub_*_31_are vectors containing the strain effects in the 24 °C and 31 °C incubations, respectively. To model the effect of the evolutionary treatment, we set, for *k ∈ {*1 *…* 29*}*:

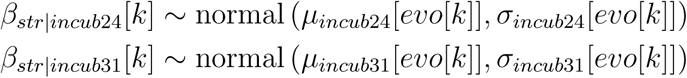

where the vector *evo* allows to map the strain ID and one of the four evolutionary treatments (three different temperature regimes plus reference strain).

The priors we used were:

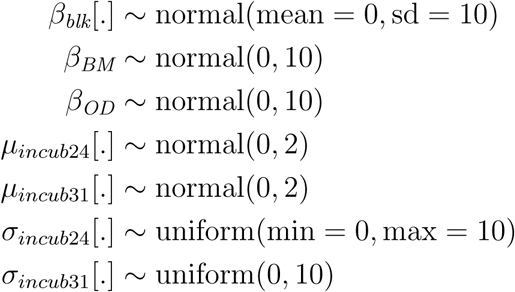

and for all *j ∈ {*1 *… T}*:

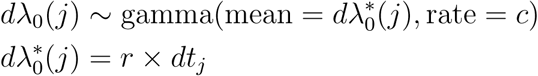

with *c* = 0.001 and *r* = 0.1. We used the first replication block and the effect of the reference strain in the 24 °C incubation as the references:

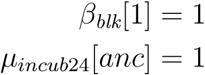

We ran four chains in parallel with the JAGS MCMC sampler for 10 000 iterations per chain, of which the first 5000 were discarded as burn-in. Model convergence and chain mixing was assessed by visual examination of trace plots and calculation of *R*^^^ values.

### 2.3 Selection of m6A in non-fully methylated GATC motifs

The method to identify GATC loci which were not fully methylated in our dataset was reported in a companion study (Bruneaux, et al., 2021). Briefly, we calculated for each GATC locus the distance between the point defined by its methylated fractions on the plus and minus strand and the point corresponding to full methylation on both strands (1,1). We then defined the set of partially methylated GATC loci of interest as the loci which deviated from the point of full methylation more than four times the average quadratic distance to (1, 1) in at least one sequenced strain.

## 3 Supplementary Tables

**Supplementary Table S1:** In-silico detection of prophage sequences in *S. marcescens* reference strain genome. Predictions were run on the PHASTER server on 2019-04-21.

| Prophage ID | Position in reference strain genome | Size (kb) | Completeness |
| --- | --- | --- | --- |
| PP1 | 521 990-535 146 | 13.2 | incomplete |
| PP4<br>(KSP20-like) | 1 970 982-2 003 867 | 32.9 | intact |
| PP5 | 3 451 823-3 468 581 | 16.8 | intact |
| PP6 | 3 914 469-3 946 913 | 32.4 | incomplete |
| PP7 | 4 423 686-4 461 768 | 34.5 | intact |

**Supplementary Table S2:**
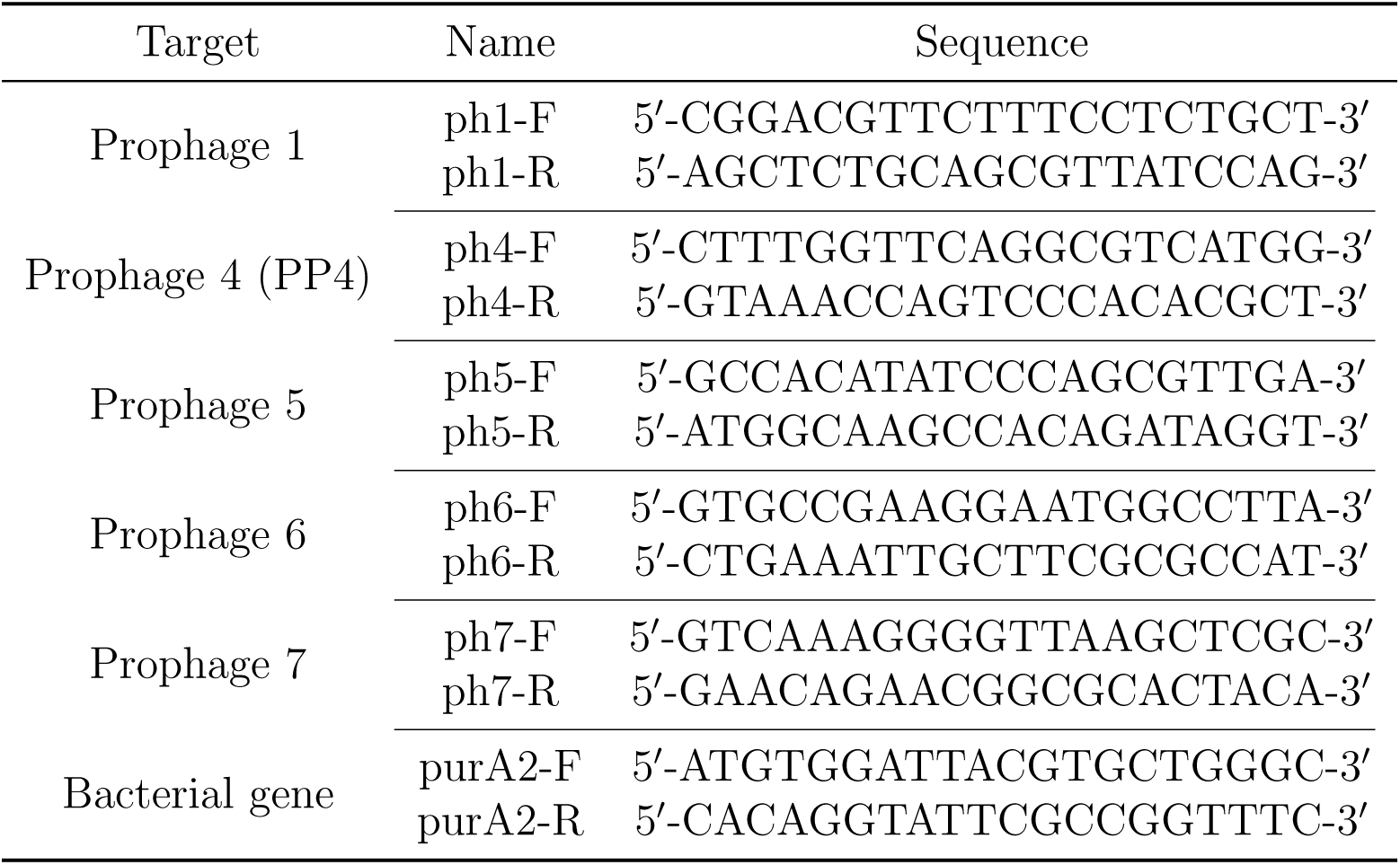
Sequences of the primers used in the qPCR quantification of prophages and chromosomal DNA. The purA2-F/R primers are targeting the chromosomal, non-prophage-related bacterial gene for adenylosuccinate synthetase.

**Supplementary Table S3:**
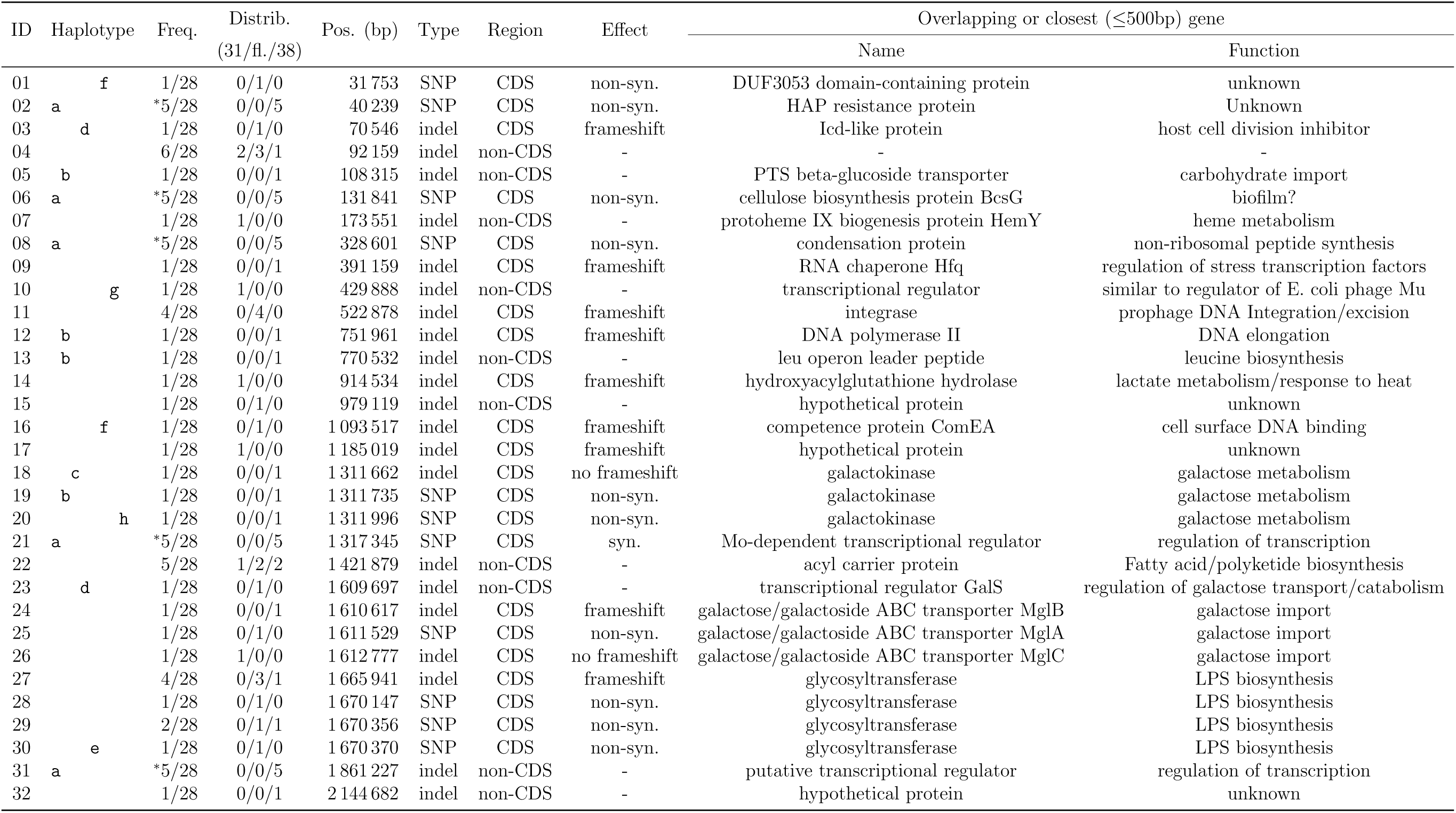
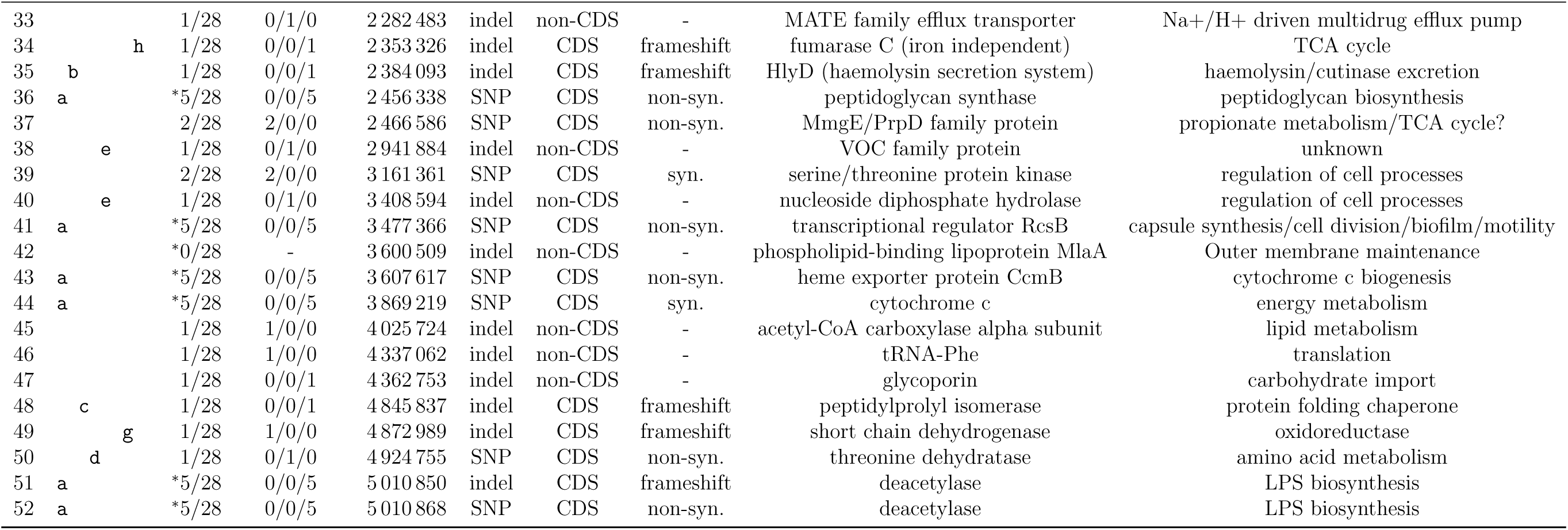
Genetic variants observed in the 29 sequenced clones (from Bruneaux et al. 2021). Haplotype: letters identify groups of co-occurring mutations. Freq.: minor allele frequency observed among the 28 evolved clones (an asterisk marks loci for which the reference strain carried the minor allele). Distrib.: distribution of minor alleles across the strains evolved in the 31 °C, 24-38 °C and 38 °C treatments.

## 4 Supplementary Figures

**Supplementary Figure S1:**
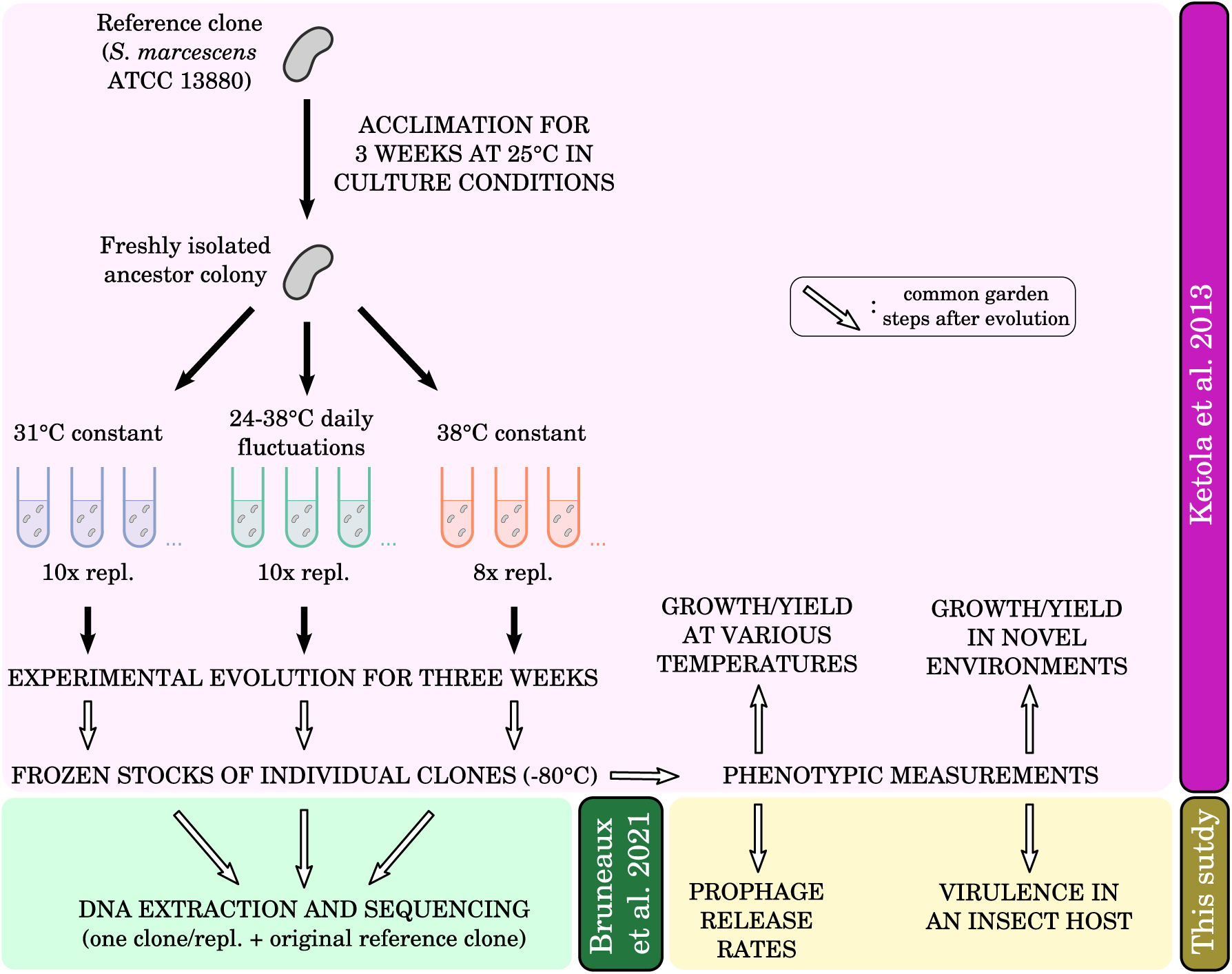
Setup of the evolution experiment from which clones were isolated and of downstream measurements. One randomly selected clone per evolved population was used for sequencing. Open arrows after experimental evolution indicates steps where evolved clones were grown under common garden conditions. Details of the evolution experiment are available in Ketola et al. (2013).

**Supplementary Figure S2:**
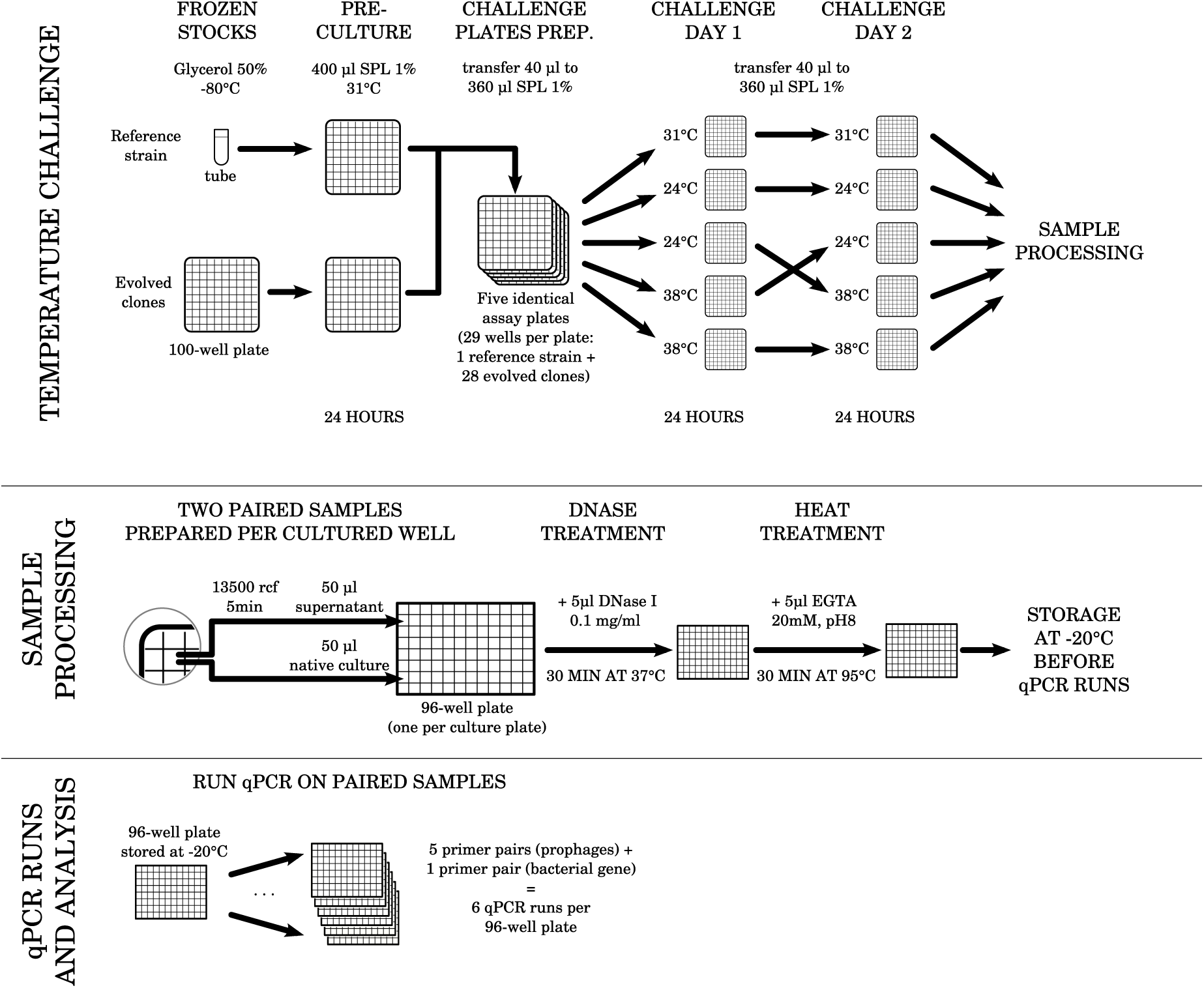
Overview of the experimental protocol used for the prophage induction assays. The prophage primers used in the qPCR runs were for prophages 1, 3, 4, 6 and 7, after preliminary experiments with the reference strain showed no detectable amount of extra-cellular DNA for prophages 2 and 5.

**Supplementary Figure S3:**
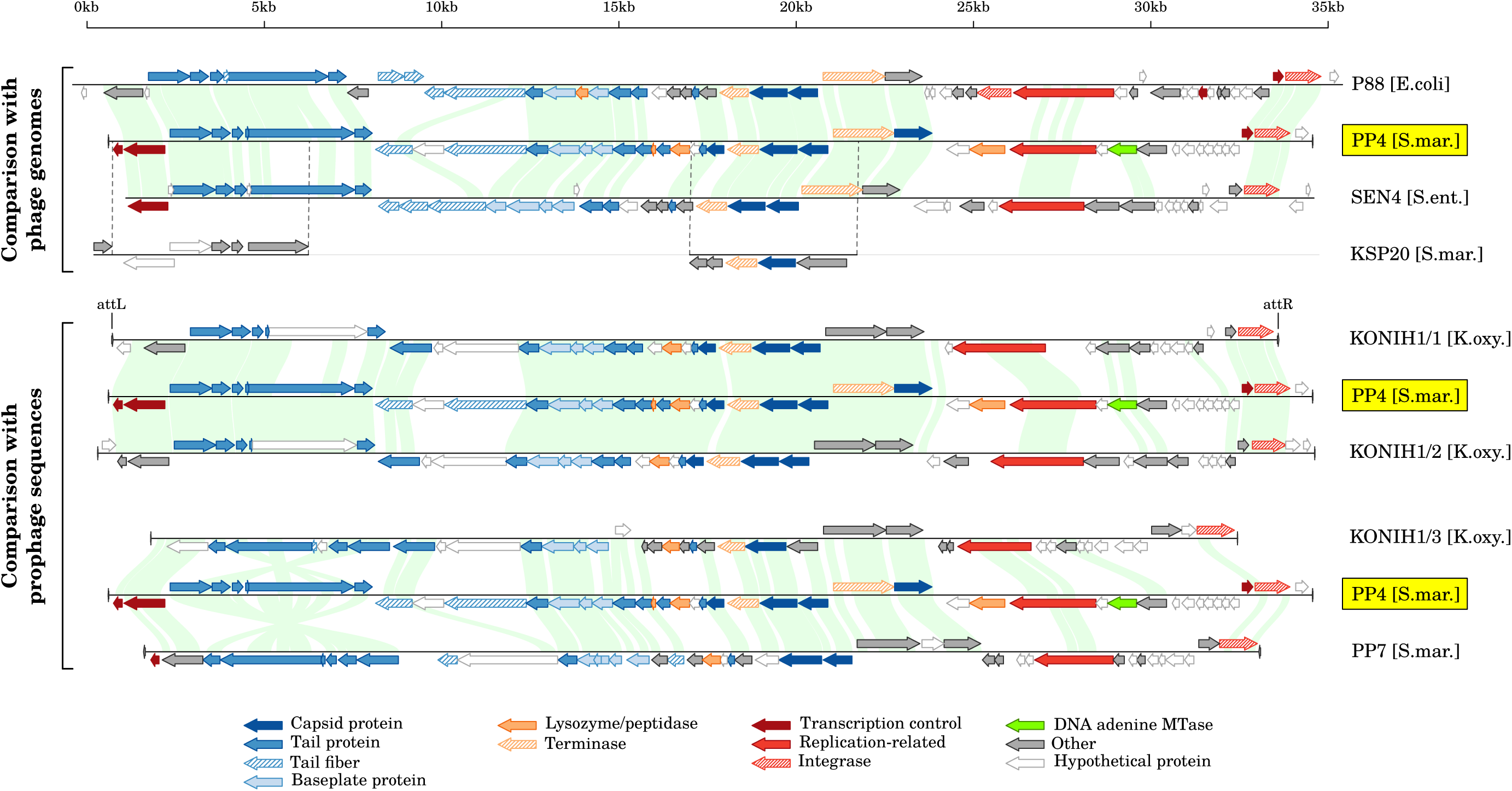
Comparison of prophage PP4 with related (pro)phages. P88 and SEN4 infect *Escherichia coli* and *Salmonella enterica*, respectively. KSP20 infects *S. marcescens* (only two sequence fragments available). KONIH1/1-3 are three prophages identified in the genome of a *Klebsiella oxytoca* strain. PP7 is another prophage candidate identified in the genome of the *S. marcescens* strain used in our study but for which no induction was detected. Matches found with `tblastx` with an e-value *≤* 10*^−^*^80^ are highlighted in green. For KSP20, those matches cover the entire areas delimited by vertical dashed lines. See details about the search for sequences related to PP4 in Methods.

**Supplementary Figure S4:**
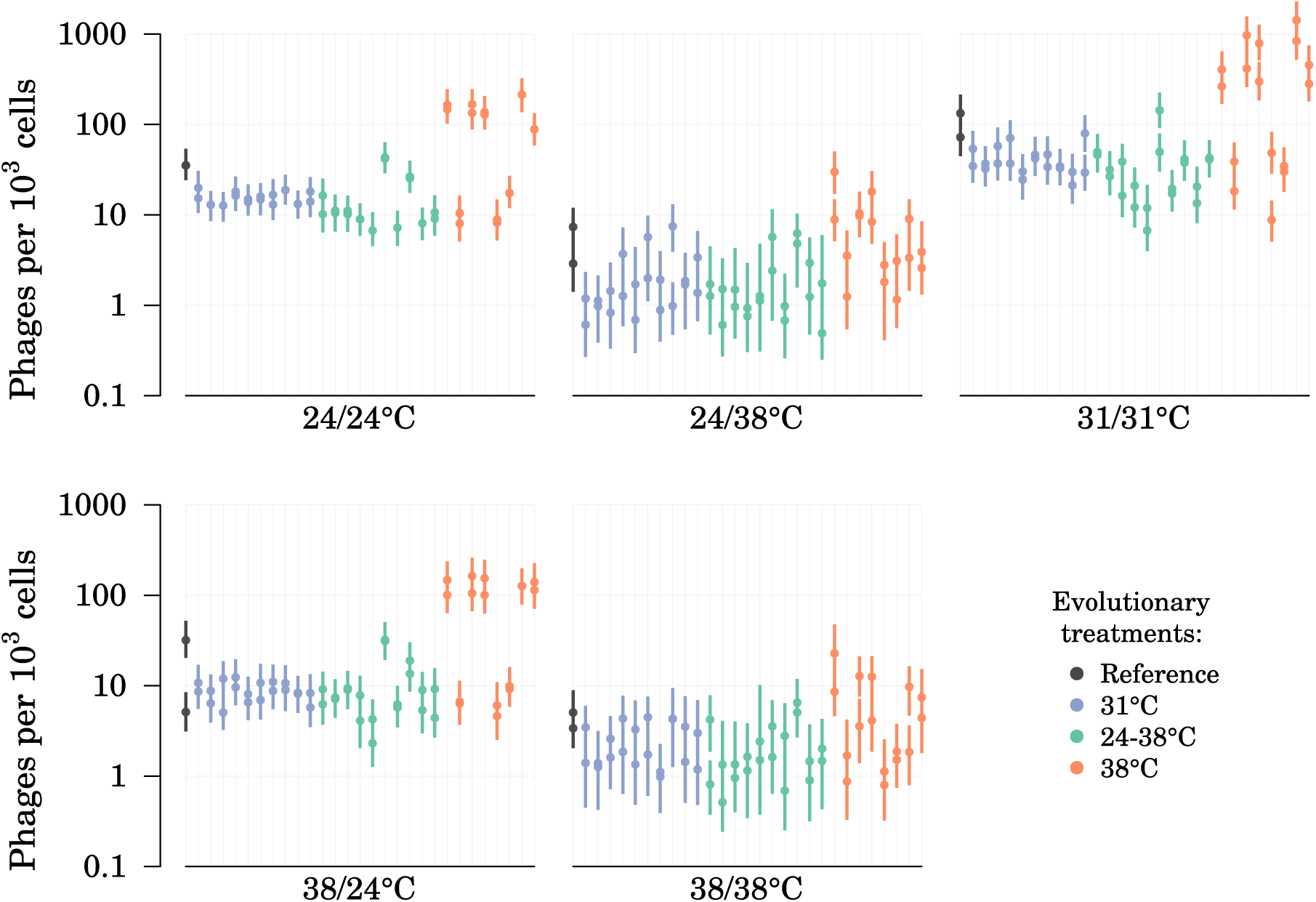
Estimated induction rates of prophage PP4 per evolved strain per assay. For each strain in each assay, two replicate measurements are available in most cases. Estimated prophage induction rates are shown as posterior mean and 95% credible interval for each measurement.

**Supplementary Figure S5:**
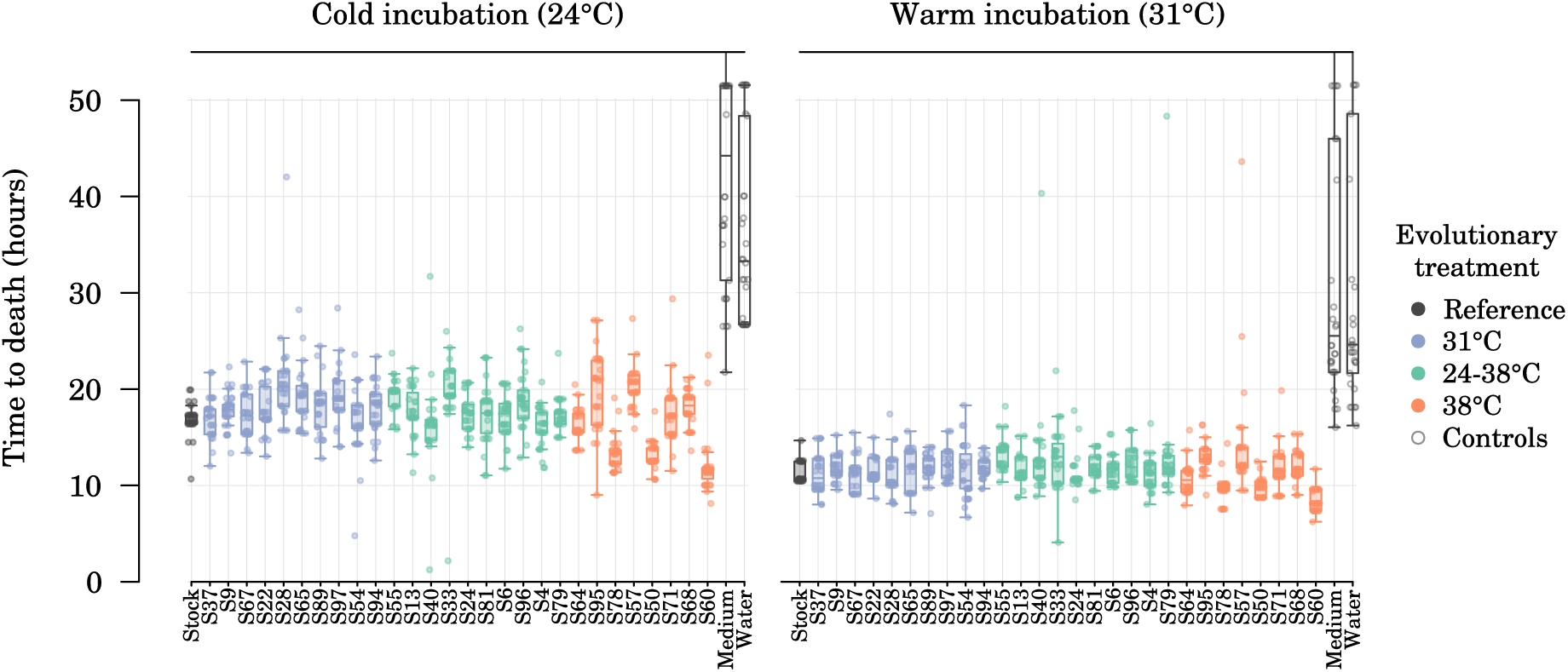
Longevity of waxmoth larvae at two incubation temperatures after injection with experimental *Serratia marcescens* strains. Each vertical lane shows larvae injected with a given strain. The Medium and Water lanes show control larvae injected with sterile medium and sterile water, respectively. Longevity is corrected for the effect of replication blocks, culture optical density and larva body mass. Dots are individual larvae. Boxplots center line, median; box limits, upper and lower quartiles; whiskers extend to the most extreme data points which are no more than 1.5 times the interquartile range away from the box.

**Supplementary Figure S6:**
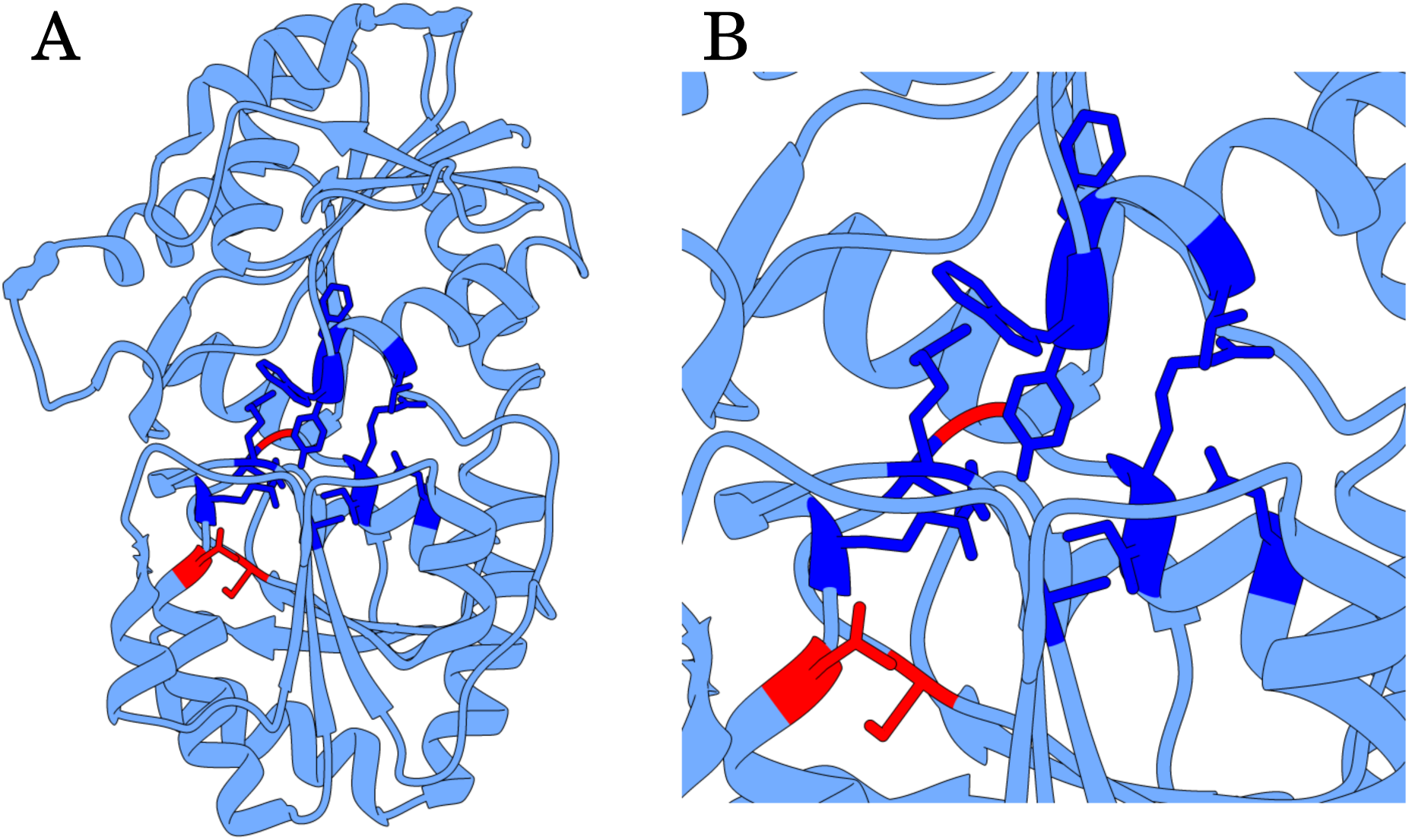
Location of observed amino acid substitutions in the predicted structure of a glycosyltransferase of *Serratia marcescens* (GenBank record QSP20947.1). The predicted protein is 380 amino-acid long and its structure was modelled using the Phyre2 online server (Kelley et al. 2015; “intense mode” run submitted on 2015-10-26). A) Overview of the tertiary structure of the complete monomer; B) close-up of the putative active site. Blue, amino-acids which have a side chain close to the ligand site, based on a protein of related structure (PDB 3mbo, Parsonage et al. 2010); red, amino-acids which exhibited three independent mutations in the evolved strains in our experiment (mutations *28*, *29* and *30* in Supplementary Table S3).

**Supplementary Figure S7:**
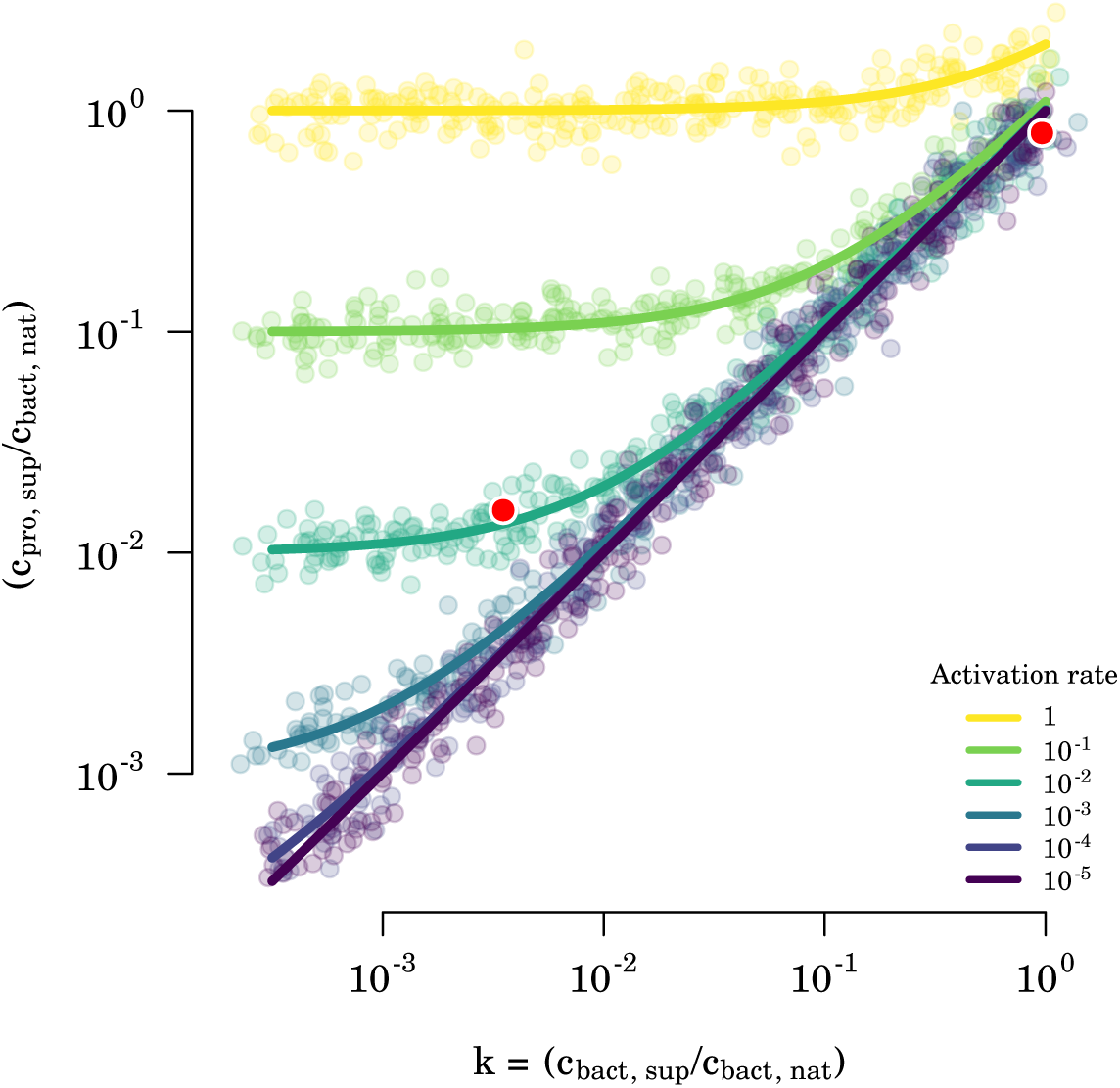
Simulation of qPCR results for different prophage activation rates *a* and different centrifugation concentration factors *k*. *c_bact,nat_*, *c_bact,sup_* and *c_pro,sup_* are the qPCR quantifications of the number of DNA copies for bacterial gene in native and supernatant samples and for prophage gene in supernatant samples, respectively. The colored lines show the predicted trajectories of *c_pro,sup_/c_bact,nat_* versus *c_bact,sup_/c_bact,nat_* from native samples (top-right corner) towards supernatant samples (to the left) as the centrifugation concentration factor *k* decreases (i.e. as supernatant samples are more and more impoverished in bacteria cells). The shape of the trajectories depends on the activation rate of the prophage, i.e. on how many phage particles are present per bacteria cells in the native sample. The colored dots matching the colored predicted trajectories represent simulations of qPCR estimations which would be obtained as the centrifugation removes more and more bacteria cells from the supernatant, assuming a precision of the Cq values *σ_cq_* = 0.48 and triplicates qPCR measurements for each culture well, as was done in our experiment. As can be seen on the figure, the sensitivity threshold to detect phage particles decreases as the depletion of bacteria cells becomes more complete. However, even at *k* values of 10*^−^*^3^, activation rates of 10*^−^*^4^ and lower are not distinguishable from the absence of induction. The red dots represent the results for a hypothetical culture, with the top-right dot representing the native sample and the bottom left dot representing the supernatant sample.

